# Alterations in retrotransposition, synaptic connectivity, and myelination implicated by transcriptomic changes following maternal immune activation in non-human primates

**DOI:** 10.1101/2020.03.31.019190

**Authors:** Nicholas F. Page, Michael Gandal, Myka Estes, Scott Cameron, Jessie Buth, Sepideh Parhami, Gokul Ramaswami, Karl Murray, David Amaral, Judy Van de Water, Cynthia M. Schumann, Cameron S. Carter, Melissa D. Bauman, A. Kimberley McAllister, Daniel H. Geschwind

## Abstract

**Background:** Maternal immune activation (MIA) is a proposed risk factor for multiple neurodevelopmental and psychiatric disorders, including schizophrenia. However, the molecular and neurobiological mechanisms through which MIA imparts risk for these disorders remain poorly understood. A recently developed nonhuman primate model of exposure to the viral mimic poly:ICLC during pregnancy shows abnormal social and repetitive behaviors and elevated striatal dopamine, a molecular hallmark of human psychosis, providing an unprecedented opportunity for mechanistic dissection.

**Methods:** We performed RNA-sequencing across four psychiatrically-relevant brain regions (prefrontal cortex, anterior cingulate, hippocampus, and primary visual cortex) from 3.5-4-year old male MIA-exposed and control offspring—an age comparable to mid adolescence in humans.

**Results:** We identify 266 unique genes differentially expressed (DE) in at least one brain region with the greatest number observed in hippocampus. Co-expression networks identified region-specific alterations in synaptic signaling and oligodendrocytes. Across regions, we observed temporal and regional differences, but transcriptomic changes were largely similar across 1^st^ or 2^nd^ trimester MIA exposures, including for the top DE genes—PIWIL2 and MGARP. In addition to PIWIL2, several other known regulators of retrotransposition, as well as endogenous transposable elements were dysregulated in MIA offspring.

**Conclusions:** Together, these results begin to elucidate the brain-level molecular mechanisms through which MIA may impart risk for psychiatric disease.

## Introduction

Epidemiological studies have implicated *in utero* environmental insults as risk factors for neurodevelopmental disorders, including schizophrenia (SCZ) (1–5). Fever and infections during pregnancy, in particular, are associated with psychiatric diagnoses in offspring, reporting adjusted odds ratios in the range of 1.2-2.5 for SCZ (4, 5) and 1.4-3.1 for autism spectrum disorders (ASD). Influenza infection as well as the pathogens that cause what are collectively called TORCH infections (toxoplasmosis, rubella, cytomegalovirus, and herpes simplex virus (HSV)) are classically known to derail CNS development in offspring when contracted during pregnancy (6, 7). Fetal exposure to toxoplasma and HSV-2 alone can as much as double the risk of SCZ in adulthood, with exposure during early pregnancy conferring the highest risk (6). These results are in accordance with twin and family studies, which have demonstrated that early childhood environmental factors contribute to liability for SCZ (8). However, despite these associations, it remains unclear how maternal infection alters brain development in offspring to increase risk for neurodevelopmental disorders. Pleiotropy between psychiatric disorders and susceptibility to infection may contribute, such that severe infections increase risk by broadly impacting early brain development, similar to other environmental risk factors (9–11). The wide range of infectious agents and exposures reported in the clinical literature suggests that it may be common factors downstream of the maternal immune response that confer risk (7).

To investigate this hypothesis, research in animal models of maternal immune activation (MIA) were developed using a poly(I:C) compound that activates the toll-like receptor 3 (TLR3) critical for recognition of many types of viruses (12). Rodent poly(I:C) MIA models have been reported to exhibit reproducible behavioral phenotypes in domains shared by those altered in human psychiatric illness, including defects in social interactions, ultrasonic vocalizations, sensorimotor gating, working memory, and cognitive flexibility, as well as elevations in anxiety and repetitive and depressive behaviors (1, 12–17). Importantly, several of these behavioral aberrations can be rescued by antipsychotics (14–16). These rodent MIA models also exhibit neuropathology and neurochemical alterations similar to those found in neurodevelopmental, psychiatric disorders (14–16, 18–24). More recently, a non-human primate (NHP) model of MIA has been developed using a derivative of poly(I:C) and their young adult offspring display abnormalities in communication, decreased social attention and interaction, and increased stereotypic behaviors that increase in intensity with age (13–16, 25, 26). NHP MIA offspring also show the molecular hallmark of psychosis—enhanced striatal dopamine, similar to the increases seen in SCZ (27). While behavioral changes in both animal models have been well characterized, the field still lacks an understanding of the molecular changes associated with MIA that may underlie these behaviors.

Transcriptomic tools such as RNA-seq enable unbiased systematic characterization of the brain level molecular changes underlying behavioral phenotypes in these models. Recently, transcriptomics has led to new causal hypotheses for SCZ and ASD related to convergence of disease risk on the regulation of synaptic genes (28, 29). In addition, elevated neural immune and inflammatory signaling pathways have been observed in postmortem transcriptomic studies across SCZ and ASD (30–35), reflecting differences in astrocyte, microglial, NFκB and interferon response pathways (29). It remains unclear, however, whether such changes could be driven by MIA and if so, when they appear during disease progression and how they relate to behavioral changes.

Here, we present the first systematic, multi-regional RNA-seq analysis of the non-human primate brain following MIA. To begin to elucidate the brain-level molecular mechanisms underlying MIA-associated behavioral alterations, transcriptomic changes were analyzed across four disease-relevant brain regions (prefrontal cortex, anterior cingulate, hippocampus, and primary visual cortex) from adolescent macaques following two *in utero* timepoints of MIA. We identify 266 unique genes differentially expressed in at least one brain region in MIA-exposed offspring. Some of these DE genes were dependent on the timing of MIA in the 1^st^ or 2^nd^ trimester, but most showed concordant changes across both exposures including the top DE genes—*PIWIL2* and *MGARP*. Our results indicate particular vulnerability of the HC to MIA and reveal altered transposable element biology, dysregulated synaptic connectivity, and enhanced myelination as important molecular pathways underlying the effects of MIA. These findings provide unique insight into the molecular changes that may precede the onset of psychosis and identify multiple new targets for mechanistic dissection *in vivo* that will help elucidate how prenatal immune insults impart risk for neuropsychiatric and neurodevelopmental disease.

## Methods and Materials

### NHP Model of MIA

All animal protocols were developed in conjunction with the California National Primate Research Center and approved by the University of California, Davis Institutional Animal Care and Use Committee. Thirteen female rhesus macaques were split into three treatment conditions: immune activation during the first trimester of pregnancy (1st Trim MIA, n=5), second trimester MIA (2nd Trim MIA, n=4), or control (Ctrl, n=4). The control group consisted of 1 untreated, 1 first trimester saline injected, and 2 second trimester saline injected NHPs. None of the injected dams in the control group showed any evidence of a maternal immune response. Animals in the MIA conditions were injected intravenously with 0.25 mg/kg of synthetic double-stranded RNA virus (poly:ICLC) (Oncovir, Inc., Washington, DC) on gestational days 43, 44, and 46 (1st Trim MIA) or 100, 101, and 103 (2nd Trim MIA).

Baseline blood samples were collected 24-48 hours before poly:ICLC injections. Additional blood samples were collected 6 hours after the second and third poly:ICLC injections. A final sample was collected approximately 5 days after poly:ICLC injections to serve as a second baseline measure. Baseline blood samples were collected during ultrasounds (animals sedated with ketamine 5-30mg/kg), injection day samples were collected without ketamine. 4ml of whole blood was collected and serum was frozen in 2ml Nalgene tubes. Serum was diluted with PBS/0.2%BSA to fall into the linear range of a primate-specific IL-6 ELISA assay (Cell Sciences, Canton, MA), and the assay was performed according to the manufacturer’s instructions (see ref 25). In the present study, remaining archived samples were processed using a non-human primate multiplexing bead immunoassay (Milipore-Sigma, Burlington, MA). The kit was run according to the manufacturer’s instructions. Briefly, 25 μL of sample was incubated with antibody-coupled fluorescent beads and then washed and incubated with biotinylated detection antibodies followed by streptavidin–phycoerythrin. The beads were then analyzed using flow-based Luminex™ 100 suspension array system (Bio-Plex 200; Bio-Rad Laboratories, Inc.). Standard curves were generated by Bio-plex Manager software to determine unknown sample concentration, with reference cytokines provided by the manufacturer in the kit. The bead sets were analyzed using a flow-based Luminex™ 100 suspension array system (Bio-Plex 200; Bio-Rad Laboratories). A five-parameter model was used to calculate final concentrations. Concentrations obtained below the sensitivity limit of detection (LOD) were calculated as LOD/2 for statistical comparisons. Supernatant aliquots were free of any previous freeze/thaw cycles.

At 3.5-4 years of age, the brain was perfused with saline, extracted, cerebellum and brainstem removed, and cerebrum bisected through the sagittal sulcus into the right and left hemispheres. The right hemisphere was sectioned with a long histology blade into ~9 6mm thick coronal slabs perpendicular to the anterior-posterior commissural (AC/PC) axis and flash-frozen in liquid nitrogen vapor. Slabs were stored in a −80 degrees Celsius freezer. Four brain regions were then dissected from the right hemisphere: the dorsolateral prefrontal cortex (DLPFC, BA 9/46) on middle frontal gyrus along the dorsal wall of the principal sulcus, anterior cingulate cortex (ACC, BA24), mid-rostrocaudal hippocampus (HC) caudal to the amygdala, and primary visual cortex (V1). For further details on animal methods, please see (25).

### RNA-sequencing and Analysis

RNA was extracted (Qiagen, miRNA easy-mini) and rRNA-depleted RNA-seq libraries were prepared using TruSeq stranded RNA plus ribozero gold kits. Libraries were multiplexed (24 per pool) and sequenced across two batches on an Illumina HiSeq2500 to an average depth of ~50 million 69 bp paired end reads per sample. Reads were mapped to the macaque genome (rheMac8) using the STAR RNA-seq aligner v2.5.0a. Gene-level quantifications were calculated with Kallisto (36) using Ensembl v87 annotations and imported into R using the tximport package. Sequence-level quality control metrics were calculated on BAM files with the PicardTools suite (CollectAlignmentSummaryMetrics, CollectRNA-seqMetrics, CollectGcBiasMetrics, MarkDuplicates). Outliers were detected by calculating standardized sample network connectivity Z scores, and samples with Z < −3 were removed from downstream analysis, as published (32). 6,410 lowly expressed genes were filtered and removed using the filterByExpr method from the R package edgeR. The filtered genes are highly enriched for olfactory receptors and ribosome related genes but not for other gene ontology categories that may be functionally important (**Figure S2A**). Samples underwent TMM normalization for read depth (37) followed by differential gene expression with Limma-voom (38). Sequencing metrics from Picard Tools were summarized by their top 4 principle components (seqPCs) (**Table S1**), which were included in the linear model to control for batch, RNA quality, and sequencing-related technical effects. As sequencing batch and RNA quality measures (sample 280:260, 260:230 ratios) were strongly correlated with these seqPCs (**Figure S1A**), they were not included in the final model as covariates. The following model was used, with specific contrast terms to determine DGE between different individual regions: *Expr ~ 0 + Group:Region + seqsPC1-4*. Differential expression p-values were then corrected for multiple comparisons using the FDRTool package in R. For select genes (e.g. *SOX10*; **Figure 7B**) that did not have an annotated homolog in NHPs, RNA-seq reads mapping to annotated human DNA sequences were used to determine expression.

The low-expressed gene filtering criteria used in the above analysis resulted in the removal of the majority of cytokine encoding genes (**Table S2**). Changes in the expression of cytokine genes are of biological interest because of previous reports of their dysregulation following MIA (39). Therefore, the differential expression of cytokine encoding genes was analyzed separately from the main dataset by fitting the same linear model above using the lmFit function in the Limma package without voom correction (**Figure S6**). Uncorrected P-values are reported for cytokine differential expression.

### Network Analysis

Network analysis was performed with the WGCNA package (40) using signed networks. A soft-threshold power of 14 was used to achieve approximate scale-free topology (R2≅0.7). Networks were constructed using the blockwiseModules function. The network dendrogram was created using average linkage hierarchical clustering of the topological overlap dissimilarity matrix (1-TOM). Modules were defined as branches of the dendrogram using the hybrid dynamic tree-cutting method (40). Modules were summarized by their first principal component (ME, module eigengene) and modules with eigengene correlations of >0.9 were merged together. Modules were defined using biweight midcorrelation (bicor), with a minimum module size of 100, deepsplit of 4, merge threshold of 0.1, and negative pamStage. Module differential expression was determined using a linear model provided by the lmFit function in the Limma package (38). Uncorrected P-values are reported for module differential expression.

### Assessment of Retrotransposition

The RepEnrich2 pipeline (https://github.com/nerettilab/RepEnrich2) was used to estimate repetitive elements in the genome using python version 2.7 (41). The repeatmasker repetitive element annotation bed file was obtained from the UCSC genome table browser (Assembly Mmul_8.0.1/rheMac8). Simple and low-complexity repeats were removed prior to running RepEnrich2. The fastq files were aligned the rheMac8 genome (Ensembl v87 annotation) using bowtie2 version 2.2.2 with settings (bowtie2 -q -p 16 -S -1 sample_R1.fastq.gz -2 sampleR2.fastq.gz output.sam -x rheMac8.fa). Samtools version 1.3.1 was used to convert sam to bam files (samtools view -bS sample.sam > sample.bam). Then RepEnrich2_subset.py (RepEnrich2_subset.py/path/to/sample.bam 30 Sample_Name --pairedend TRUE) was run on the output bam files to subset uniquely mapping and multimapping reads. RepEnrich2.py (RepEnrich2.py /data/rheMac8_repeatmasker.bed/data/Sample_Output_Folder Sample_Name /data/RepEnrich2_setup_rheMac8/data/sample_name_multimap_R1.fastq --fastqfile2 /data/sample_name_multimap_R2.fastq/data/sample_name_unique.bam --is_bed TRUE --cpus 16 --pairedend TRUE) was used to obtain count files for the different repetitive elements. Significant repetitive elements were determined in the same way as DGE, as described above, using Limma-voom (38).

### Enrichment Analyses

Over-representation analysis among the top up and downregulated gene sets and for co-expression network modules was performed using a one-sided Fisher’s exact test on predetermined gene lists with relevance to neurodevelopmental or psychiatric disorders. The SFARI gene list was defined as the list of high-confidence category 1 and 2 ASD risk genes (https://www.sfari.org/resource/sfari-gene/). The SCZ and ASD up and down regulated genes lists were obtained from the PsychEncode consortium (29). The p-value cutoff used for up and downregulated gene list was 1e-5 for SCZ, 0.01 for ASD, and 0.05 for all genes showing suggestive association with MIA in this study. P-values from over representation analysis were Bonferroni corrected for multiple comparisons. Over-representation analyses between co-expression networks identified in this paper and those identified in PsychEncode (29) and Winden et al. (42) were performed using the same approach as above. Multiple testing correction was performed across all pairwise comparisons between NHP MIA modules and the modules reported in both previous analyses and specifically reported for mitochondrial enriched modules of interest (**Figure S9C, D**).

g:ProfileR (43) was used to determine gene ontology enrichment in GO:bp, GO:cc, GO:mf, and KEGG among the top up and downregulated gene sets with a p-value cutoff of 0.05 and for co-expression network modules. An ordered query was performed by the signed P-value for differentially expressed gene lists and by kME for co-expression network modules. Both analyses used *M.mulatta* related terms with strong hierarchical filtering, FDR correction with a maximum set size of 1000 genes.

Expression Weighted Cell-type Enrichment (EWCE) (44) was used for cell-type enrichment analysis among the top up and downregulated gene sets with a p-value cutoff of 0.05 and for co-expression network modules. Cell-type specificity for the expression of each gene was determined using human brain single-nucleus RNA-seq (Nuc-seq) data from the PsychEncode Consortium and Lake et al., 2018 (45). FDR corrected p-values less than 0.05 are reported as significantly enriched.

### Rank-Rank Hypergeometric Overlap (RRHO)

Original gene lists from this experiment and from studies of post-mortem SCZ and ASD brains were ordered by signed -log10 P-value and compared via a one-sided hypergeometric test with a step size of 100 followed by Benjamini-Yekutieli FDR correction using the *RRHO* package in R (46). Final p-values are reported on a −log10 scale.

### Western Blotting

Frozen Brain tissue (HC or DLPFC) was disrupted by sonication with a probe sonicator in 2x SDS Lamelli buffer (15μl buffer/mg brain tissue) and immediately denatured at 85°C for 5 minutes. Samples were then centrifuged at 10,000 RPM for 10 minutes to remove any non-soluble debris from the lysate. The concentration of protein in the sonicated lysates was determined by a BCA protein assay (Thermo). DTT was added to a final concentration of 100mM. 15ug of protein per sample was loaded and run on a 4-15% TGX precast gel (Biorad). Proteins were then transferred to methanol activated PVDF membranes for 90 minutes at 100V. Membranes were then blocked in LI-COR TBS blocking buffer (LI-COR Biosiences), and probed overnight or for 3 hours with primary antibodies, blots were than washed with TBS tween (0.05% tween) and probed with secondary antibodies matched to the host species of the primary antibody and conjugated to fluorescent dyes (1:15:000, LI-COR). Blots were then washed and imaged on an Odyssey CLX imager (LI-COR). Primary antibodies used were mouse anti-GAPDH (1:10,000, Sigma, Cat # G8795), sheep anti-PIWIL2 (1:500, R&D Biosciences, Cat # AF6558), rabbit anti-CX3CR1 (1:2000, Sigma, Cat # C8354), rabbit anti IGSF6 (1:250, Atlas Antibodies, Cat # HPA041072), Bands were quantified in the Image Studio software and probe bands were normalized to GAPDH band signal to control for protein load. Differences between groups were determined by performing 2 tailed *t*-tests in GraphPad Prism software.

## Results

### Maternal immune activation induces brain transcriptomic changes in adolescent NHPs

RNA-sequencing was performed on 4 brain regions to characterize brain transcriptomic changes in a well characterized cohort of 3.5-4 year old male non-human primates (NHPs; **Figure 1**). NHPs were exposed *in utero* to saline or poly:ICLC on 3 days starting at gestational day GD44 (1^st^ Trimester MIA) or GD101 (2^nd^ Trimester MIA; **Methods and Materials**). Following quality control (**Figures 1D, S1, S2A**), differential expression (DE) was characterized within and across the brain regions profiled (DLPFC, ACC, HC, and V1) for all 14,975 brain expressed genes detected (**Methods**).

**Figure 1.**
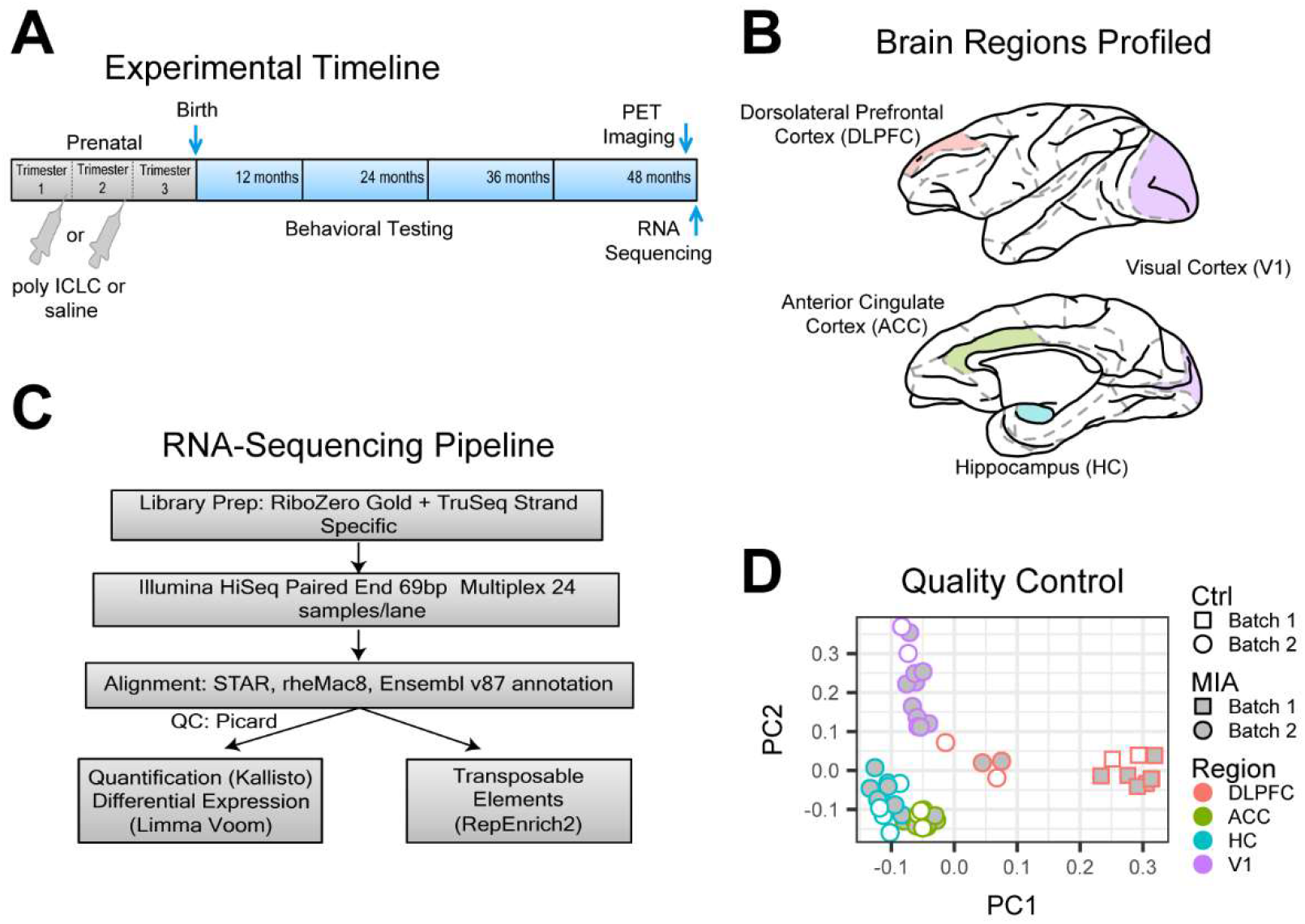
Outline of experimental approach and bioinformatic analyses. **(A)** Female non-human primates were injected with the viral mimic Poly:ICLC at either gestational day GD44 (1st trimester) or GD 101 (2nd trimester). Tissue was harvested from male offspring at 48 months followed by RNA-sequencing. Behavioral testing for stereotypic behaviors was performed throughout development and PET imaging was performed directly before sequencing and has been previously published on this same cohort (26) **(B)** Four brain regions with relevance to schizophrenia and autism biology were profiled including dorsolateral prefrontal cortex (DLPFC), cingulate cortex (ACC), hippocampus (HC), and primary visual cortex (V1). **(C)** Diagram of RNA-sequencing pipeline. Briefly, RiboZero stand specific library preparation was performed before obtaining 69bp paired end lllumina reads. Reads were aligned to the current version of the Rhesus genome and quantified with Kallisto or RepEnrich2, which is specially designed to detect the expression of transposable elements (TEs). In both cases differential expression was quantified using a linear mixed effects model with Limma-voom. **(D)** Top principal components (PCs) of normalized gene expression demonstrating that batch and brain region are the largest drivers of variation in the dataset.

Six genes showed globally significant patterns of differential expression at FDR < 0.1 -- *PIWIL2*, *MGARP*, *C15orf41*, *SNED1*, *FCRL3*, *RNASE1* (**Figure 2A**; **Table S2**). The top DE gene, *PIWIL2*, is a master regulator of piRNA mediated DNA methylation and an inhibitor of retrotransposition (47, 48). *PIWIL2* also regulates the expression of plasticity-related genes in the adult brain, and disruption of the piwi pathway in HC enhances contextual fear memory in mice (49). The second highest DE gene, *MGARP,* encodes a protein found on mitochondria that regulates mitochondrial morphology, distribution, and motility. Decreased MGARP expression in neurons causes dendritic and axonal overgrowth in mice (50). The other globally significant genes are not well characterized in regulating brain development or function.

**Figure 2.**
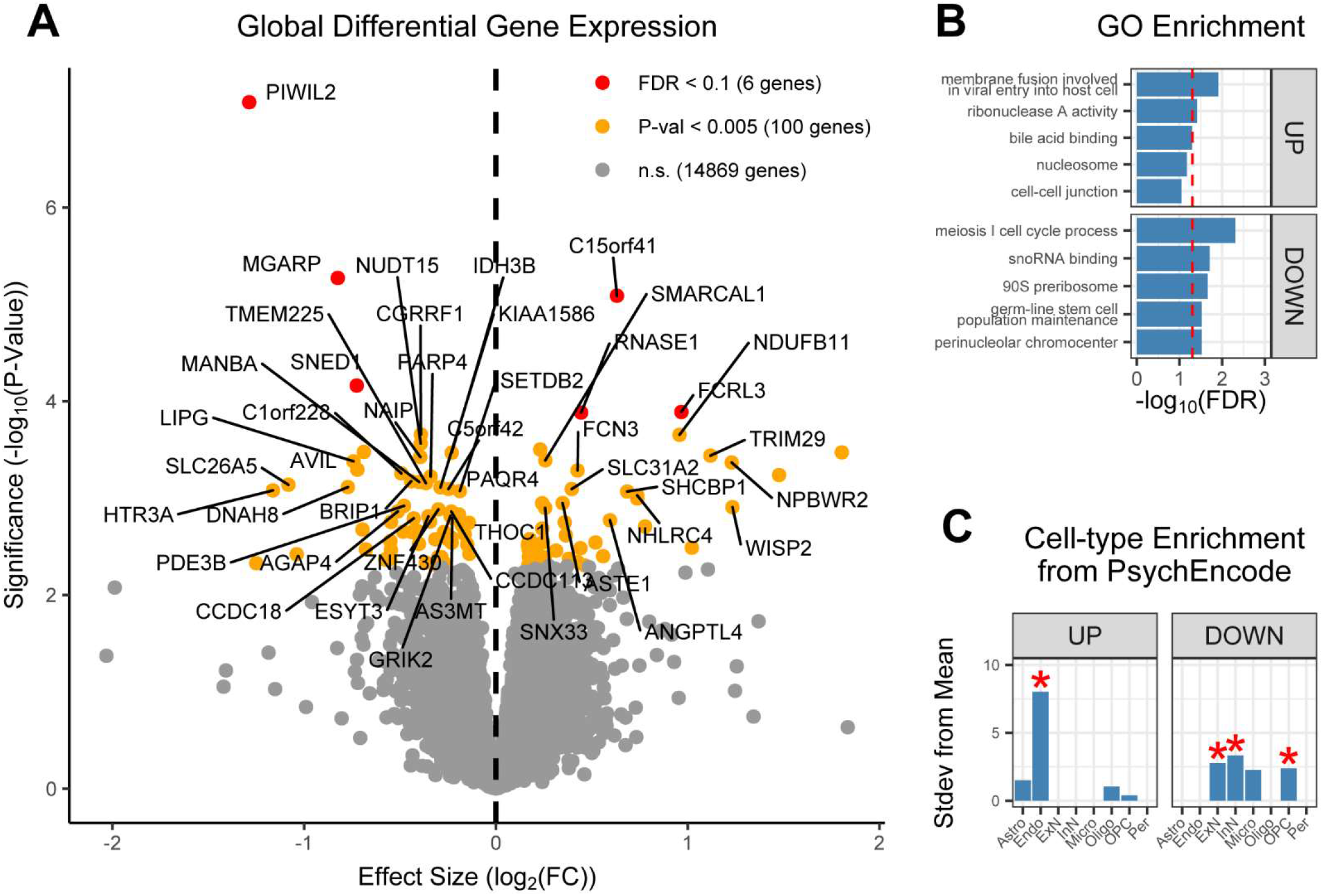
Global differential gene expression across brain regions following MIA. **(A)** Differential gene expression analysis using Limma-voom was performed on all saline injected vs. poly:ICLC injected MIA samples and pooled across brain regions and MIA timepoints. Volcano plot of differential gene expression detects PIWIL2 as the top globally downregulated gene followed by MGARP. Red dots indicate genes that pass FDR correction for differential gene expression (FDR<0.1), yellow dots indicate suggestive association with MIA, and grey dots indicated minimal or no association. Dotted line indicates log2(FC)=0 Genes are labelled according to their human homologs, those without annotated homologs are left unlabeled. **(B)** Top GO terms enriched among up and downregulated genes with p<0.05 following MIA determined using g:ProfileR. Red dotted line indicates an FDR significance threshold of 0.05. **(C)** Cell-type specificity of up and downregulated genes following MIA with p<0.05 genes based on PsychEncode and Lake et al, 2018 adult human single-cell Nuc-seq data. *FDR<0.05, unlabeled cell-types are not significantly enriched.

An additional 100 genes showed suggestive global association with MIA (P < 0.005), including several downregulated neurotransmitter receptors (*HTR3A*, *GRIK2*, *GLRA2*) and the high-confidence SCZ risk gene *AS3MT* (51). Gene ontology enrichment analyses identified significantly up and downregulated terms including *membrane fusion involved in viral entry into host cell* with core enrichment of *HYAL2*, *GAS6*, and *NECTIN2* (**Figure 2B**). In addition to its role in viral entry, *GAS6* promotes oligodendrogenesis and myelination in the adult nervous system (52) and acts as a neurotrophic factor during hippocampal development (53). *NECTIN2* is necessary for neuronal and glial survival (54). Cell-type enrichment analyses using human single-nucleus RNA-seq data from PsychEncode (45) was performed using EWCE (44). Upregulated genes (P < 0.05) showed enrichment for endothelial cell markers, whereas excitatory neuron (ExN), inhibitory neuron (InN), and oligodendrocyte precursor cell (OPC) markers were enriched among downregulated genes (**Figure 2C**). These findings demonstrate that MIA-induced changes in brain gene expression affect disease relevant genes, cell-types, and pathways that persist into adolescence in this model.

### Specificity of brain molecular changes based on prenatal timing of MIA

In rodent MIA models, the timing of immune activation during pregnancy specifies the nature and severity of neurodevelopmental and behavioral outcomes in affected offspring (1, 17). Here, we tested the role of MIA timing on gene expression in the brains of offspring by comparing two susceptibility windows in NHPs whose gestation closely parallels that in humans (55). MIA timing-specific DE was characterized separately for NHPs exposed to 1^st^ or 2^nd^ trimester MIA (**Figure 3A**; **Table S3**). None of the control animals injected with saline at either time-point or non-injected had measurable immune activation and so, they were pooled for these comparisons. Overall, transcriptomic changes were largely concordant across timepoints (*R^2^* of log_2_-fold changes of 0.54, P<10^−15^) (**Figure 3B**) taking the limited sample size into consideration. *PIWIL2* and *MGARP* again emerged as the top downregulated genes during each time point independently. In contrast, several genes showed more specific patterns of DE. For example, the serotonin receptor *HTR3A* was only significantly downregulated following 1st trimester exposure. In contrast, 2nd trimester exposure showed unique upregulation of several genes, including *IFITM3. IFITM3* has been shown to mediate the brain immune response to neonatal poly(I:C) exposure, disrupts intracellular steroid metabolism, and is strongly upregulated in post-mortem brains from subjects with SCZ (33, 56, 57). The 2nd trimester signal also showed selective enrichment of RNA processing related gene ontology pathways among downregulated transcripts, indicating that some differences do exist (**Figure 3C, D**). However, given that the global signatures showed highly significantly concordance, we grouped the timepoints together for downstream analyses so as to boost power.

**Figure 3.**
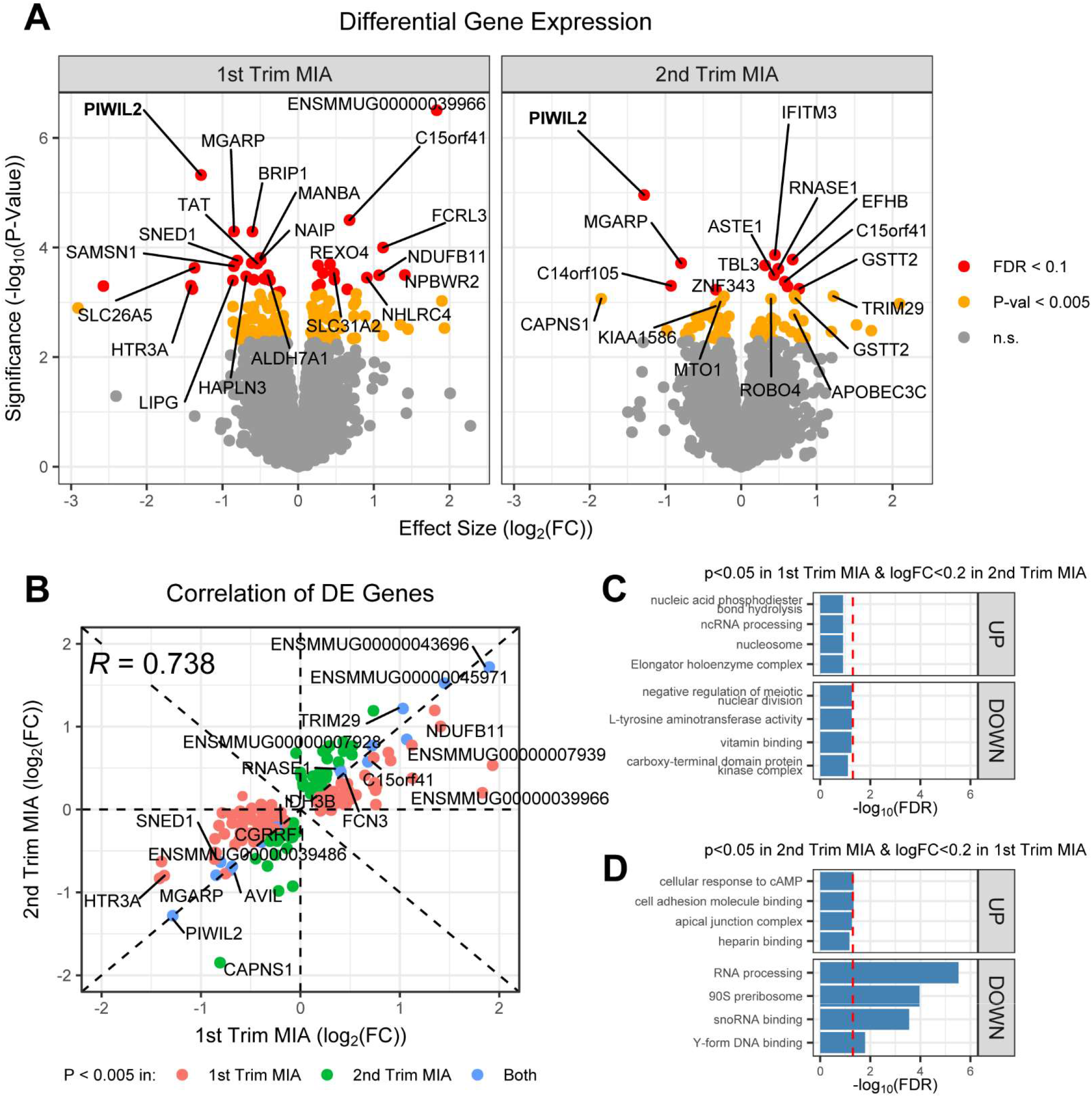
High concordance between genes differentially expressed in first and second trimester MIA offspring. **(A)** Differential gene expression analysis using Limma-voom was performed separately on saline injected vs. poly:ICLC injected MIA samples from 1st or 2nd trimester pooled across brain regions. Volcano plots of differential gene expression detect PIWIL2 and MGARP as the top downregulated genes following both 1st and 2nd trimester MIA. Red dots indicate genes that pass FDR correction for differential gene expression (FDR<0.1), yellow dots indicate suggestive association with MIA, and grey dots indicated minimal or no association. Genes are labelled according to their human homologs, those without annotated homologs are left unlabeled. **(B)** All genes with suggestive association with MIA p<0.005 following either timepoint of poiy:ICLC injection were plotted comparing their log2(FC) following either 1st or 2nd trimester MIA. Blue dots indicate genes that are associated with MIA regardless of timepoint while red and green dots indicate specific dysregulation following 1st or 2nd trimester MIA respectively. The overall log2(FC) correlation between genes dysregulated following either timepoint is R=0.738. **(C, D)** To determine the trimester specific effects of MIA, GO enrichment was determined for genes with p<0.05 following 1st trimester MIA and log2(FC) <0.2 following 2nd trimester MIA and vise-versa using g:ProfileR. Red dotted line indicates an FDR significance threshold of 0.05.

### MIA alters gene expression in the brains of offspring in a region-specific manner

Although there is region- and temporal-specificity in alterations in cytokine protein levels in the brains of offspring following MIA (39), whether there is region-specificity in gene expression remains unknown. To address this, we compared patterns of DE across 4 distinct brain regions, including cortical (DLPFC and V1) and limbic regions (ACC and HC) (**Figure 4A**). The greatest number of DE genes was observed in HC (n=118 genes; FDR<0.1), followed by V1 (n=114) and DLPFC (n=34), with only minor DE detected in ACC (**Figures 4A, B**; **Table S2**). Again, *PIWIL2* emerged as a downregulated gene across 3 of the 4 regions (DLPFC, ACC, and HC). Focusing on individual DE genes, however, may fail to detect small yet concordant transcriptomic changes shared across regions. To investigate this possibility, we compared log_2_-FC effect sizes for all genes exhibiting suggestive association (P<0.005) with MIA across regions. Similarities were low, but significant (p < 1e-15), with the largest correlation (*R* = 0.447) between the DLPFC and ACC and other regions (DLPFC and V1; *R* = 0.121; p = 0.00397) appearing more distinct (**Figure 4C**). These associations were smaller or non-existent when comparing the concordance in log_2_-FC across all genes (**Figure S2B**). Together, results clearly show that MIA causes DE of genes in a region-specific manner in offspring.

**Figure 4.**
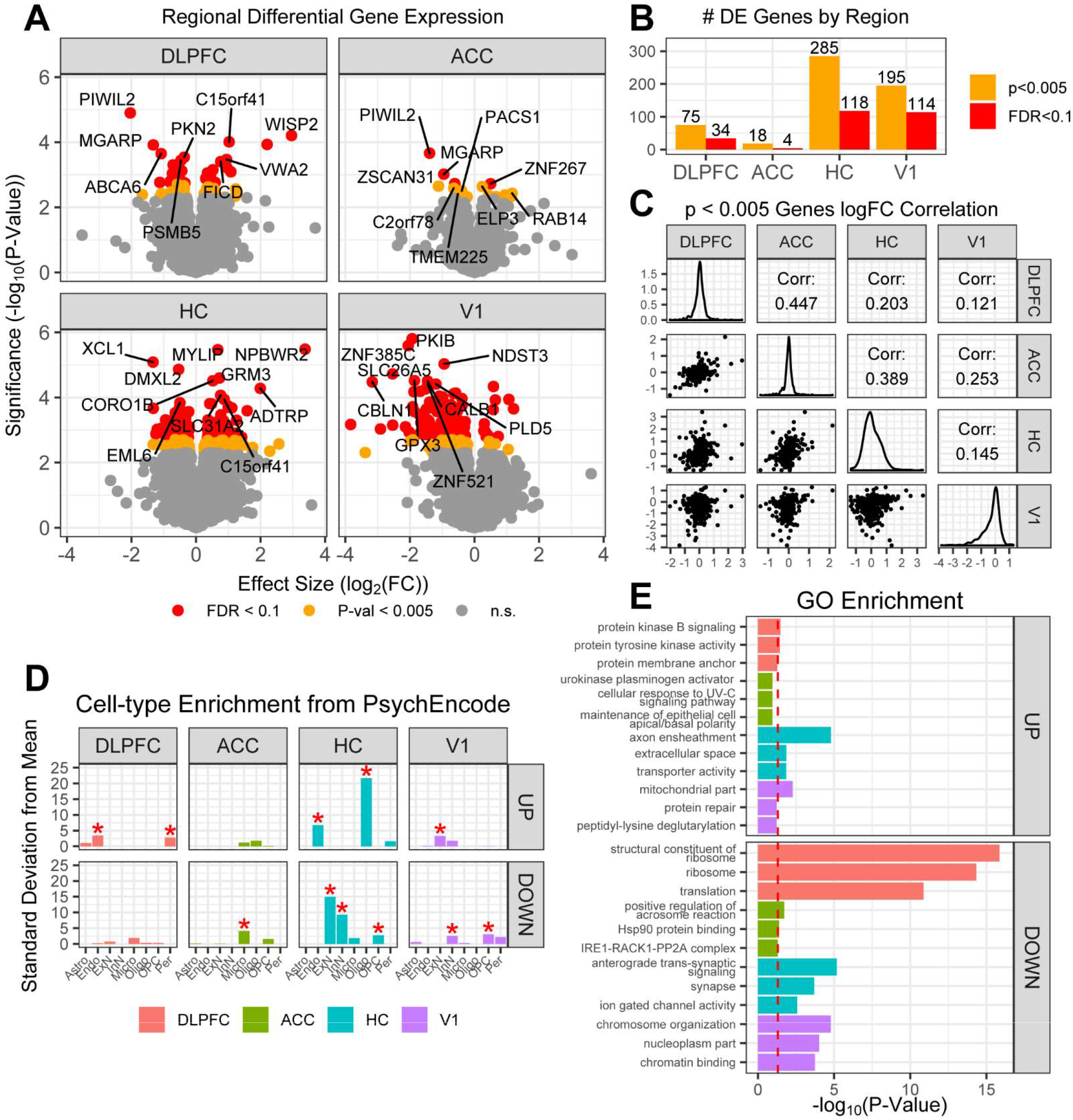
MIA alters gene expression in a region-specific manner in the brains of offspring, **(A)** Differential gene expression analysis using Limma-voom was performed separately for each brain region on all saline injected vs poly:ICLC injected MIA samples pooled across MIA timepoints. Volcano plots indicate the top 10 differentially expressed in each region following MIA. Red dots indicate genes that pass FDR correction for differential gene expression (FDR<0.1), yellow dots indicate suggestive association with MIA, and grey dots indicated minimal or no association. Genes are labelled according to their human homoiogs, those without annotated homologs are left unlabeled **(B)** The number of DE genes in each region by significance threshold. **(C)** log2(FC) off all genes with suggestive association (p<0.005) with MIA in at least one brain region are compared pairwise with each other brain region. Small correlations indicated minimal overlap in the similarity of gene expression changes between each region following MIA. **(D)** Celi-type specificity of region specific up and downregulated genes following MIA with p<0.05 genes based on PsychEncode and Lake et al, 2018 adult human single-cell Nuc-seq data. *FDR<0.05, unlabeled cell-types are not significantly enriched. **(E)** Top GO terms enriched among region specific up and downregulated genes with p<0.05 following MIA determined using g:ProfileR. Red dotted line indicates an FDR significance threshold of 0.05.

The region-specific effects of MIA also segregate by cell-type and biological process. Specifically, upregulated genes in the HC are strongly enriched in oligodendrocytes, whereas downregulated genes are strongly enriched for ExNs, InNs, and OPCs (**Figure 4D**). Interestingly, the ACC is the only region that contains any cell-type enrichment for microglia among downregulated genes (**Figure 4D**). Region specific differences in cell-type enrichment are also concordant with GO term enrichment determined using g:ProfileR. The enrichment of upregulated “axon ensheathment” and downregulated “anterograde trans-synaptic signaling” related genes closely matches cell-type enrichment results observed in the HC and is again specific to that region (**Figure 4E**). Meanwhile, the strongest GO enrichment is for changes in “structural constituent of the ribosome” among downregulated genes in the DLPFC. This change is specific to the DLPFC and indicates that changes in mRNA translation may also be relevant in maternal immune activation models. Enrichment among downregulated genes in V1 is also specific for “chromosome organization” (**Figure 4E**).

### Convergent gene expression changes following MIA implicate transposable elements and steroid biology

To more specifically capture the molecular processes underlying transcriptomic changes, we next performed weighted gene correlation network analysis (WGCNA) across all samples, detecting 29 gene co-expression modules after iteration across multiple network parameters (**Figure 5A**; **Figure S2C-E**; **Table S4**). The steelblue module was downregulated across all brain regions with an effect size (log_2_FC = −0.074) nearly double the next most significantly changing module (**Figure 5B, C**). It contained 105 genes of which 41 lack human homologs. Surprisingly, many of these 41 genes appeared to be heavily associated with transposable element (TE) biology. 8/41 belong to the “LINE-1 RETROTRANSPOSABLE ELEMENT ORF2 PROTEIN” family (PTHR25952:SF231; ref (58)), including module hub genes *ENSMMUG00000041596* and *ENSMMUG00000039238*. Other hub genes include *HERV-H LTR-Associating 2* (*HHLA2*), an immune checkpoint molecule, and *Tigger Transposable Element Derived 1* (*TIGD1*), a paralog of centromere binding protein *CENPB* (**Figure 5D**). Both of these genes were previously TEs that have been evolutionarily co-opted into protein coding genes (59). Gene ontology enrichments were observed for pathways related to steroid/lipid metabolism including *INSIG2*, *SULT2A1*, and *FAAH2* (**Figure 5E**). Notably, only MEsteelblue and only one other module -- MEyellow -- had the same direction of change across all brain regions analyzed while all others experienced divergent expression (**Figure 5F**).

**Figure 5.**
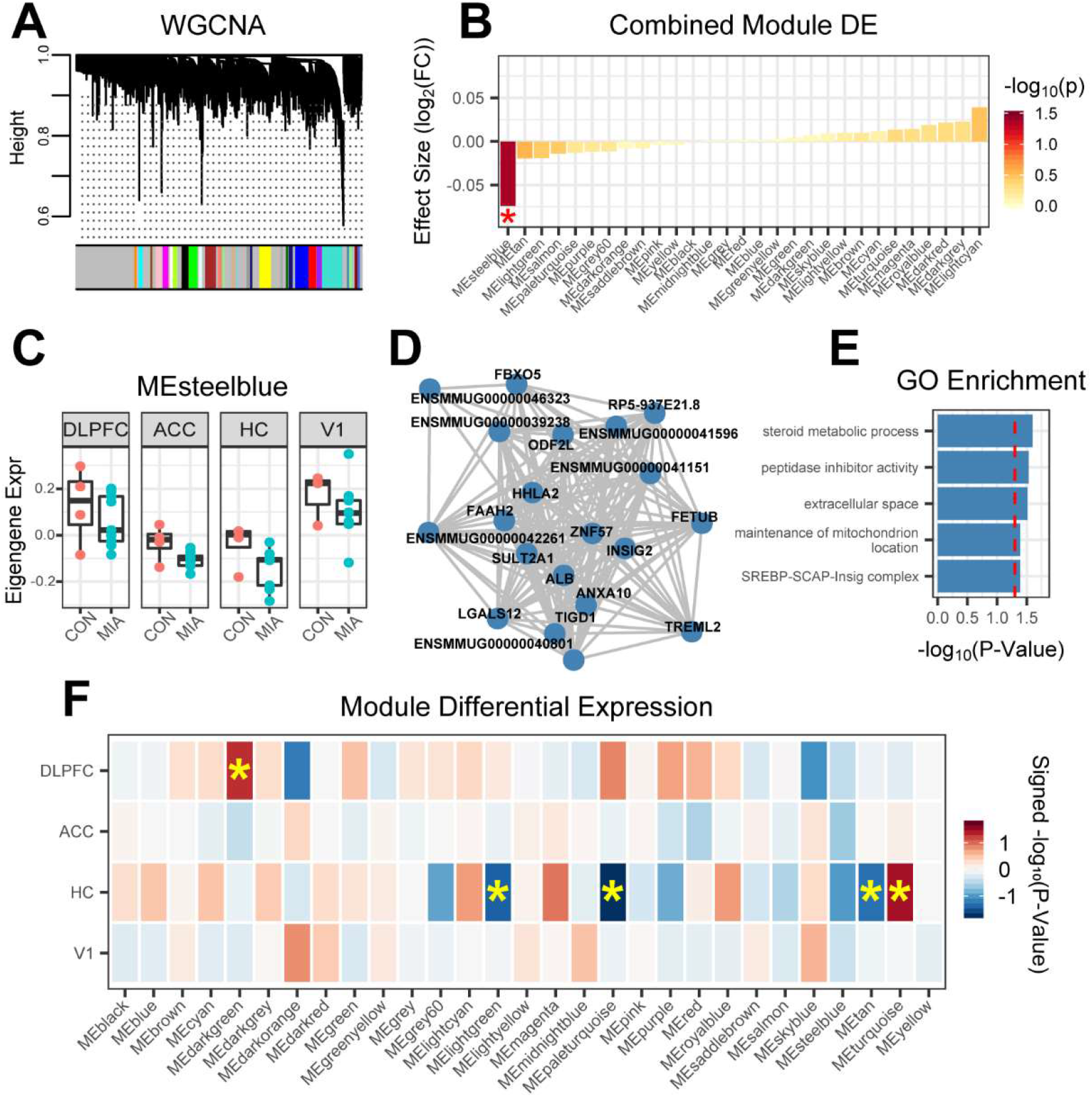
Consistent changes across brain regions in MIA offspring implicates transposable elements and steroid biology. **(A)** WGCNA was used to construct a signed bicor network and identify co-expression network modules, each containing genes whose expression is highly correlated across samples. Dendrotree height indicates the degree of co-expression correlation and colors indicate co-expression network module assignments. **(B)** Differential co-expression network expression was determined using Limma-lmFiton all saline injected vs. poly:ICLC injected MIA samples pooled across brain regions and MIAtimepoints. Only one co-expression network, MEsteelblue, is significantly downregulated across all pooled samples. *p<0 05, **(C)** Boxplot of MEsteelblue module eigengene expression across the brain regions analyzed, **(D)** Top 20 hub genes for MEsteelblue. **(E)** Top GO terms enriched in MEsteelblue region by g:ProfileR. For all GO enrichment plots, red dotted line indicates an FDR significance threshold of 0 05 **(F)** Differential co-expression network expression was determined using Limma-lmFit separately for each brain region on all saline injected vs. poly:ICLC injected MIA samples pooled across MIAtimepoints. *p<0.05.

Notably, the top DE gene across all brain regions, *PIWIL2*, is a master regulator of piRNA mediated DNA methylation and an inhibitor of retrotransposition (47, 48). After confirming that *PIWIL2* is also downregulated at the protein-level (**Figure 6A-C**), we sought to determine whether specific classes of TEs are more broadly dysregulated following MIA. Expression of TEs was quantified from RNA-seq reads using RepEnrich2 (41), a computational tool for the genome-wide estimation of TE abundance (**Figure S3A-C**). Only one specific TE, *HERV1_LTRa*, was detected with an FDR < 0.1 in the ACC while a total of 26 separate TEs had suggestive association with MIA (p < 0.005) in at least one brain region. These include multiple elements of the human endogenous retrovirus (HERV) and long terminal repeat (LTR) families of retrotransposons which have been previously shown to contain transcription factor binding sites and can modulate transcription (60). Alterations in specific LINE-1 elements L1M6 and L1M3de were also observed in the DLPFC (**Figure 6D, S3D**). Other alterations include region specific DE of piggy-bac derived PGBD3 and PGBD5 which have been co-opted into protein coding genes (59) (**Figure S3E, F**). Together, these results indicate specific changes in transposable element biology similar to what has been observed in other neurological diseases as a response to inflammatory signals (61, 62).

**Figure 6.**
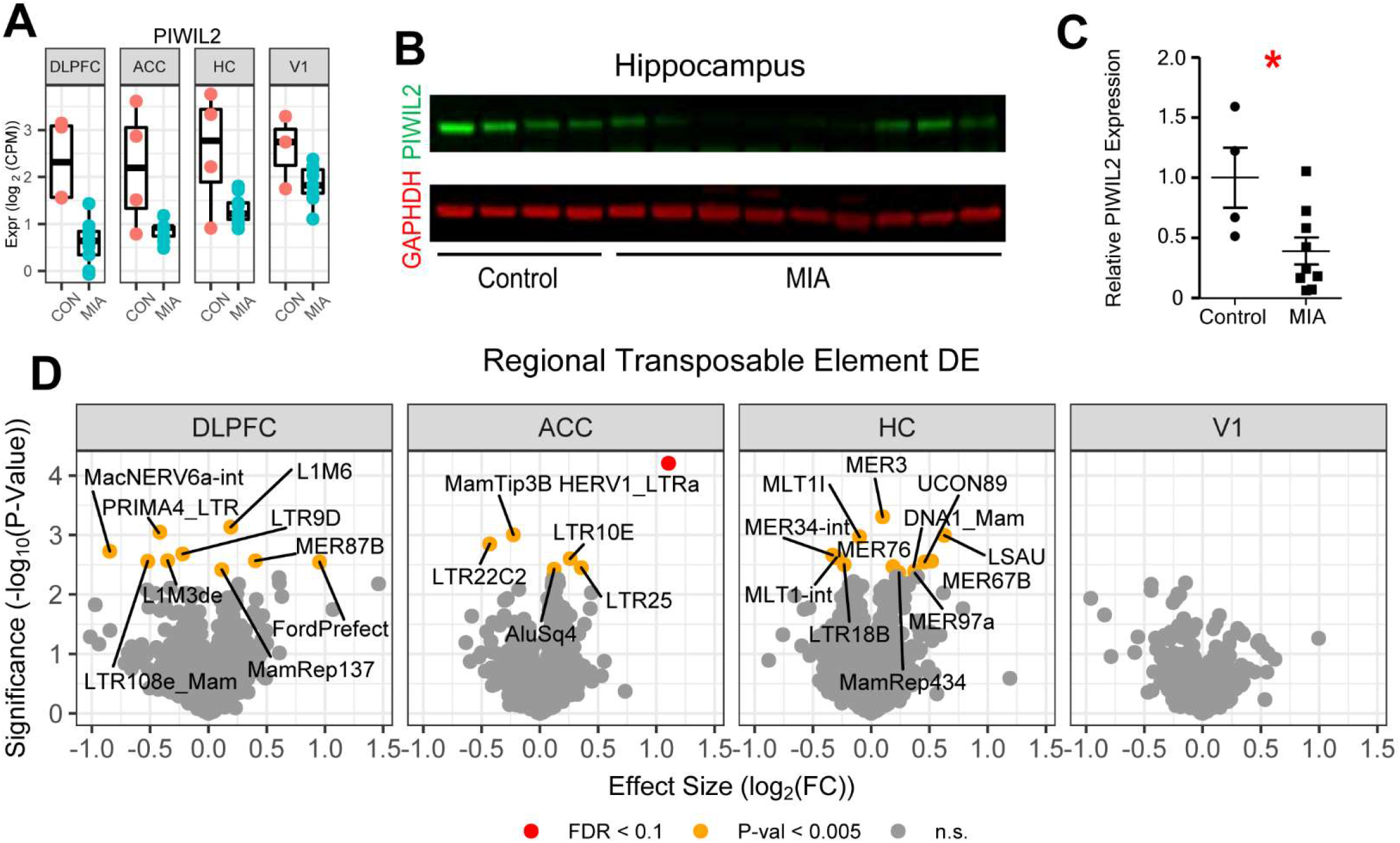
MIA causes dysregulation of transposable elements across brain regions in offspring. **(A)** Boxplot of piRNA regulator PIWIL2 expression across brain regions following MIA. **(B)** Western blot of PIWIL2 expression on the hippocampus following MIA. **(C)** Quantification of part **B** relative to GAPDH. *p<0.05. **(D)** Differential transposable element expression analysis using Limma-voom was performed separately for each brain region on all saline injected vs. poly:ICLC injected MIA samples pooled across MIA timepoints. Volcano plots indicate all transposable elements with suggestive association with MIA in each brain region, Red dots indicate transposable elements that pass FDR correction for differential transposable elements expression (FDR<0.1), yellow dots indicate suggestive association with MIA, and grey dots indicated minimal or no association.

### Hippocampal co-expression networks implicate synaptic downregulation and increased myelination

Several modules were most specifically dysregulated in HC, often with divergent patterns in other regions. Indeed, we detected both the largest number of differentially expressed genes (**Figure 4A, B**) and co-expression modules (**Figure 5F**) in HC, suggesting greater biological vulnerability to the effects of MIA in this region. Upregulated genes in HC showed enrichment for oligodendrocyte and endothelial cell markers, whereas downregulated genes were enriched for excitatory neurons, inhibitory neurons, and oligodendrocyte precursor cells (OPCs) (**Figure 4D, E**). These results are broadly concordant with GO term enrichment, finding “axon ensheathment” as the most strongly enriched term among upregulated genes and “anterograde trans-synaptic signaling” as most strongly enriched among downregulated genes (**Figure 4D, E**). Synaptic genes included a number of glutamate receptors and high-confidence autism risk genes including upregulation of *GRM3* and *GRM8*, and downregulation of *TRIO* (**Table S2**). Additionally, upregulation was detected among all genes defining a core regulatory network important for the differentiation of oligodendrocytes and the production of myelin sheaths including *OLIG2*, *SOX10*, *MYRF*, *PLLP*, *MBP*, *CNP*, *MOG*, and *MAG* (63) (**Figure 7A, B**), suggesting a hyper-myelination phenotype in the HC.

**Figure 7.**
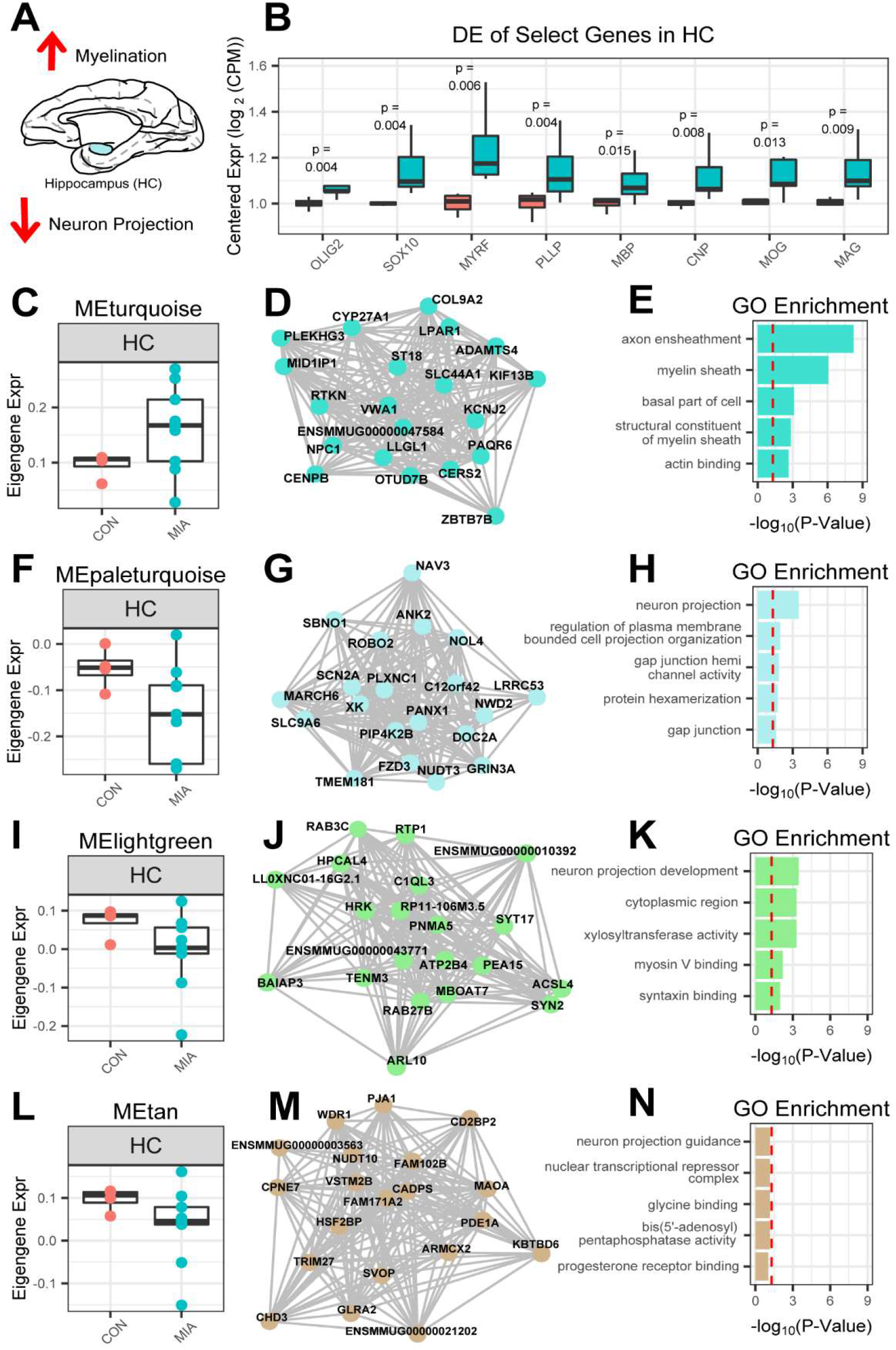
MIA increases myelination and oligodendrocyte-related genes and decreases neuron projection genes in hippocampus of offspring. **(A)** Summary of changes in the HC. **(B)** Multiple oligodendrocyte and myelin sheath related genes are upregulated in the HC following MIA. **(C)** Boxplot of MEturquoise module eigengene expression across the brain regions analyzed. **(D)** Top 20 hub genes for MEturquoise. **(E)** Top GO terms enriched in MEturquoise region by g:ProfileR. **(F)** Boxplot of MEpaleturquoise module eigengene expression across the brain regions analyzed. **(G)** Top 20 hub genes for MEpaleturquoise. **(H)** Top GO terms enriched in MEpaleturquoise region by g:ProfileR. **(I)** Boxplot of MEIightgreen module eigengene expression across the brain regions analyzed. **(J)** Top 20 hub genes for MEIightgreen. **(K)** Top GO terms enriched in MEIightgreen region by g:ProfileR. **(L)** Boxplot of MEtan module eigengene expression across the brain regions analyzed. **(M)** Top 20 hub genes for MEtan. **(N)** Top GO terms enriched in MEtan region by g:ProfiieR. For all GO enrichment plots, red dotted line indicates an FDR significance threshold of 0.05.

These findings were similarly observed among HC co-expression network modules, with the most significantly up- and downregulated modules, MEturquoise and Mepaleturquoise (**Figure 7C, F**), enriched for genes related to axon ensheathment and neuron projections, respectively (**Figure 7D-E, G-H**). Other notable genes in MEturquoise include *NPC1* and *NPC2* which regulate cholesterol export from lysosomes and are non-functional 19 in Neimann-Pick Type C disease (64). MEpaleturquoise on the other hand contains high-confidence autism risk genes *SCN2A* and *ANK2* among its hubs and multiple ion channels including *NAV3* and *SLC9A6* (**Figure 7G**). Other dysregulated co-expression network modules in the HC include MElightgreen which is enriched for “neuron projection development” (**Figure 7I-K**) and MEtan which contains no significant GO term enrichment (**Figure 7L-N**). MElightgreen counts among its hub genes multiple members of the Ras family of G-protein coupled receptors *RAB3C* and *RAB27B* that have known roles in calcium mediated synaptic vesicle release (65, 66) (**Figure 7J**). Meanwhile MEtan is notable for containing the hub gene *CADPS*, another high confidence autism risk gene whose drosophila ortholog functions in calcium mediated synaptic vesicle release and whose loss disturbs glutamatergic neurotransmission (67) (**Figure 7M**).

### Minimal immune signature in MIA NHP brains

Because neuroinflammatory gene expression changes have been reported in SCZ and ASD (29) and because cytokines are altered in a region- and temporal-specific manner in the mouse MIA brain throughout postnatal development (39), we next examined whether there were modules enriched in glial and immune genes in the NHP MIA brains. Although we detected co-expression modules exhibiting strong cell-type enrichment for astrocyte (MEgreen) and microglial (MEblue) markers, differential expression of these modules was not observed within or across brain regions (**Figures 5B, F, S4**). An orthologous analysis of cell-type specific marker gene enrichment among nominally DE genes (P<0.05) showed similar results, with the exception of a possible slight decrease in microglial cell markers in ACC (**Figure 4D**). The lack of a major microglial signature is consistent with recent reports from the MIA mouse model indicating little microglial dysregulation in the brain of adult MIA offspring (68) and from the human literature showing conflicting results from PET imaging studies (69).

While it appears that MIA NHP brains lack a broad immune signature, specific examples of immune dysregulation do exist. Although gene expression of the majority of cytokines whose protein expression was previous analyzed in a mouse model of MIA (39) is unchanged in the NHP MIA brains, specific upregulation of CCL3, CXCL1, and IL-5 is detected (**Figure S6**). Notably, the expression changes in these cytokines are brain region specific, with CCL3 and IL-5 solely upregulated in HC while CXCL1 is specifically upregulated in V1. Additional genes of interest include microglial markers *CX3CR1* and *IGSF6* of which only *IGSF6* is downregulated and shows suggestive association (P<0.005) with MIA in the DLPFC (**Figure S7A, B**). These mRNA expression results contrast with protein level data from western blot in both DLPFC and HC. Particularly, while *CX3CR1* trends upward in the DLPFC and downward in the HC of MIA offspring, its protein levels are significantly down-regulated in the DLPFC with no change in HC. *IGSF6* is trending downwards in both DLPFC and HC, but its protein levels are significantly upregulated in both regions (**Figure S7C, D**). Overall, these results suggest compensatory mechanisms counteracting each other at the level of transcription or mRNA translation and suggest that there may be other protein level immune related changes that are missed by our current transcriptomic analysis.

### Limited overlap in gene expression changes between adolescent NHP MIA offspring and adult SCZ and ASD samples

When accounting for the estimated conversion of 1 macaque year to 4 human years (55), the non-human primates examined here (3.5-4 NHP years; 14-16 human equivalent years) are significantly younger than human subjects included in the genetic and postmortem analysis for SCZ (median age > 30; ref (29)). Thus, results from the NHP model may provide unique information about the molecular changes that precede the onset of psychosis. To determine if the molecular signature in the MIA NHP model at this young age overlaps with the published signatures found in older humans with SCZ or ASD, we performed a rank-rank hypergeometric overlap test (RRHO; ref (46)) on genes ranked by their degree of differential expression in MIA vs their DE rank in ASD or SCZ as assessed in DLPFC by PsychENCODE (**Figure S8A**). No broad or region-specific overlap was detected between the ranked genes from SCZ or ASD samples and from our study -- even when focusing on DLPFC, which is most represented among samples from the PsychEncode Consortium (29) (**Figures S8B**). However, downregulated genes from two regions -- HC and V1 -- showed slight enrichment for high-confidence ASD risk genes (SFARI) (**Figure S8C**) and individual DE genes in multiple brain regions have been linked to ASD or SCZ, as indicated above. Although we cannot determine if MIA ultimately causes similar transcriptomic changes in adult NHP offspring, as in humans with ASD or SCZ, the signatures in our adolescent offspring indicate molecular changes that are likely critical for the progression of the MIA-induced alterations and could provide insight into changes that might relate to a prodromal period in humans (98).

We also sought to determine whether any gene expression signatures exhibited associations with the increased, repetitive stereotyped behaviors observed in MIA exposed animals (**Figure 8A**; **Table S5**). The total number of stereotypies is weakly correlated with module eigengene expression across regions, with the strongest correlations detected in V1. MEmidnightblue and MEdarkorange in particular have the highest correlations with the number of stereotypies (log2 transformed; *R* = 0.64, P < 0.027; *R* = 0.63, P < 0.027, respectively) (**Figure 8B, S9A, S9B**). MEmidnightblue displays significant GO enrichment for mitochondrial membrane compartment while MEdarkorange is enriched for mitochondrial compartment, suggesting that cellular respiration in the visual cortex may be loosely associated with repetitive behaviors (**Figure 8C-F**). These modules also display statistically significant overlap with mitochondria enriched module geneM33 from PsychEncode (29) (**Figure S9C**). Previous work from our lab has also identified mitochondria specific co-expression networks that are differentially enriched in synaptic and non-synaptic cellular compartments (42). We observe a nominally significant overlap between the non-synaptic (cell body) mitochondrial co-expression module from Winden et al. (42) and MEmidnightblue, suggesting that the modules identified in this paper may also be related to activity dependent energetic function (**Figure S9D**).

**Figure 8.**
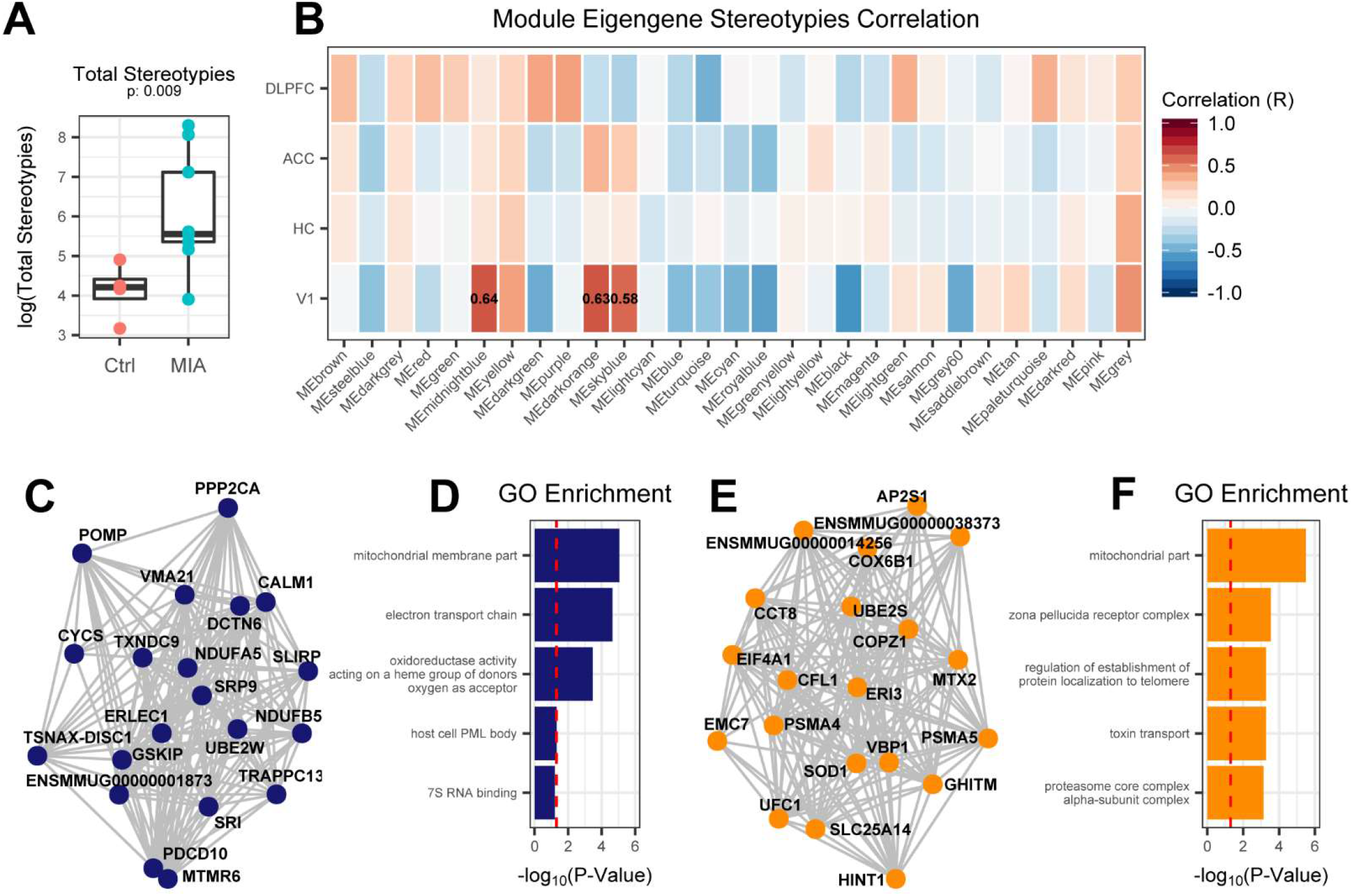
Mitochondrial co-expression networks in V1 correlate with behavioral aberrations in offspring. **(A)** The total number of stereotypic behaviors was measured in NHPs throughout development. The log2(Total # stereotypies) observed is significantly increased in MIA offspring as compared to controls (p=0.009; Student’s t-test). **(B)** Correlation between module eigengene expression in a given region and the log2(Total # stereotypies) in a given subject. The largest correlations with log2(Total # stereotypies) are found in V1, particularly MEmidnightblue and MEdarkorange. **(C)** Top 20 hub genes for MEmidnightblue **(D)** Top GO terms enriched in MEmidnightblue by g:ProfileR. **(E)** Top 20 hub genes for MEdarkorange (F) Top GO terms enriched in MEdarkorange by g:ProfileR. For all GO enrichment plots, red dotted line indicates an FDR significance threshold of 0.05.

Finally, we examined whether the severity of maternal interleukin 6 (IL-6) induction following poly:ICLC exposure is correlated with brain molecular phenotypes in our NHP data (**Methods; Table S5**). MEdarkorange and MEsalmon in particular have the highest correlation with the measured maternal IL-6 levels (*R* = 0.65; *R* = 0.64 respectively) (**Figure S9E**). As noted above, MEdarkorange is highly correlated with increased stereotypies and is enriched for genes in the mitochondrial compartment. MEsalmon displays significant GO enrichment for “regulation of Ras protein signal transduction” although none of the co-expression networks that correlate with maternal IL-6 are differentially expressed in any region.

## Discussion

Here, we provide the first transcriptomic analysis across multiple brain regions of a recently developed NHP model of MIA. We observe robust pan-cortical and unique regional signatures of differential gene expression. Our findings implicate alterations in transposable element biology, synaptic connectivity, and myelination with relative hippocampal vulnerability in the adolescent NHP brain following MIA. Although we find no significant overlap between this NHP model and patterns of DE observed in adult patients with SCZ and ASD, these NHPs are significantly younger than human subjects included in genetic and postmortem analysis for ASD and SCZ. Moreover, analysis of the functional pathways implicated by our analysis in the MIA NHP brains are associated with the pathophysiology of both disorders and the magnitude of changes in several co-expression modules correlate with aberrant behaviors. We hypothesize that the transcriptional changes reported here may provide unique insight into prodromal changes in the brain that precede psychosis (98).

MIA has been hypothesized to exert its effect through epigenetic changes which then present dynamically over the trajectory of brain development and maturation (1, 7). Along these lines, multiple studies have reported changes in DNA and histone methylation following MIA (70–73). The most strongly downregulated gene in our dataset, *PIWIL2*, which is detected at the mRNA and protein level, is also known to act as a global regulator of DNA methylation through its interactions with piRNAs (47, 48). Thus, if it is correct that MIA primarily exerts its effect through epigenetic changes, *PIWIL2* may act as a driver of downstream changes that are specific to each brain region.

Alternatively, *PIWIL2* may induce changes in gene expression in a secondary manner, by altering regulation of transposable elements, which have shown signs of increased activation in previously published MIA models (72, 74). In this regard, since *PIWIL2* plays a role in transposon silencing (47, 48), a consistent increase in TE expression would be expected in brain regions in which it is depleted. Instead, we observe both up and down-regulated TEs with suggestive association with MIA (**Figure 6D, S3D**) and a downregulated module, MEsteelblue, with an unexpectedly large number of predicted LINE-1 elements (**Figure 5B-D**). Changes in transposable elements have been associated with both SCZ (75) and ASD (76), but it is unclear whether and how the complex changes in our NHP MIA dataset are related. These data also challenge the simple hypothesis that *PIWIL2* affects gene expression only through the downstream effects on transposable elements, and suggests the possibility that *PIWIL2* also may regulate the expression of downstream genes through methylation changes in MIA models. Indeed, in the adult HC, it has recently been reported that *PIWIL2* regulates the expression of plasticity-related genes with disruption of the piwi pathway enhancing contextual fear memory and causing hyperactivity in mice (49).

One of the strongest effects of MIA in the brains of adolescent NHPs is altered transcription of genes related to myelination and found in oligodendrocytes. Consistent with this result, defects in myelination have been reported in other MIA models, as well as in human disease (77, 78). Myelination deficits were detected in the PFC, HC, and nucleus accumbens (NAc) in adolescent offspring from the MIA mouse model (70, 79), although region specific findings differed between studies. qRT-PCR for multiple neuronal, glial, and inflammatory markers in a pig model of MIA conversely detected a strong *increase* in *MBP* expression in addition to a decrease in neuron density in the HC at fetal timepoints (80). This result is broadly in line with the increase in myelination and decrease in neuron projection-related genes observed in adolescent NHPs in this study, suggesting that some MIA related outcomes are either not detectable in rodents or are detectable at different ages than were examined. All of these studies are limited in ease of comparison due to differences in the timing of MIA exposure. Since altered myelination-related genes have also been reported in both ASD and SCZ (81, 82), our results highlight the importance of future experiments to determine the progression of MIA-induced changes in myelination through development to adulthood and the causal relationship of these changes to the neuropathological and behavioral phenotypes in offspring.

Previous MIA studies have shown reproducible behavioral changes in MIA mouse models (83, 84), which have led to a search for a common biological underpinning. Transcriptomic and proteomic studies have examined both embryonic (85–87) and adult (69, 88, 89) timepoints and long-term changes were related to G-protein coupled receptor signaling and glutamatergic and serotonergic receptors, similar to our NHP model. Changes in synaptic connectivity have been hypothesized to be a potential point of convergence for the many diverse genes and biological processes dysregulated in ASD (90, 91). MEdarkgreen, which is downregulated in the DLPFC contains *CNTNAP2*, which causes a severe recessive neurodevelopmental syndrome associated with hyperactivity, language dysfunction and ASD (92), as one of its hubs. Additionally, multiple co-expression networks in the hippocampus including MEpaleturquoise, MElightgreen, and MEtan contain genes known to localize in dendritic spines. Especially striking is that all of these co-expression networks demonstrate down-regulation, suggesting a very broad dysregulation at the synapse.

Non-synaptic MIA-induced DE genes may also converge on this common pathway to alter neural connectivity. Consistent with this idea, the two most highly DE genes in the NHP MIA brain, *PIWIL2 and MGARP,* alter expression of plasticity-related genes and dendritic and axonal arborization in mice (49–50). Another example is brain-derived neurotrophic factor (BDNF), which regulates dendritic arborization, connectivity, and plasticity during typical brain development (93) and is decreased in the aged brain following MIA (68), BDNF expression itself trends downward in the HC (log2-FC = −1.06, p = 0.03) and BDNF is included in the MElightgreen module which is significantly downregulated in the HC and is enriched for genes with the function of regulating neurotransmitter levels (**Figure 7J, S4B**). Since BDNF has previously been implicated in the regulation of myelination (94), it represents a potential unifying link between the diverse changes observed following MIA.

Mitochondria enriched co-expression networks MEmidnightblue and MEdarkorange may also provide insights into the behavioral phenotypes in our NHP model. Particularly, both modules from visual cortex (V1) correlate with the number of stereotypies observed in NHPs. There is no known relationship between V1 and repetitive behaviors, so this may reflect more widespread alterations that were simply detectable in this region, perhaps because of its high neuronal density (95). Repetitive behaviors are a hallmark of multiple neurodevelopmental and neuropsychiatric illnesses and further assessment of other brain regions that have been associated with repetitive behaviors, such as striatum is warranted in future studies (96, 97).

Although future work is clearly needed to determine if MIA ultimately leads to similar molecular changes in adult NHP MIA offspring as in humans with SCZ, our results are relevant for understanding how MIA leads to molecular phenotypes and behavioral aberrations in adolescent offspring in disease-related domains. Importantly, our data may reveal molecular changes prior to the onset of psychosis, that are not accessible in human samples. In this regard, the differences in transcriptomic changes in the MIA and human disease datasets are potentially consistent with results from the mouse model indicating that MIA causes dynamic and brain region-specific changes in molecules that are age-dependent, with the opposite direction of change for cytokine protein levels found in adolescence compared to young adulthood (39). Alternatively, MIA may be just one component of disease risk that needs to be combined with genetic or other environmental factors to lead to conditions such as ASD or SCZ.

Despite the inherent limitation of limited sample sizes available when working with NHPs and the limitation of transcriptomic approaches to uncover functional protein level changes, the functional pathways implicated by our analysis in the MIA NHP brains have been implicated in SCZ and therefore are worthy of deeper investigation. Particularly interesting lines of future work will focus on the role of *PIWIL2* methylation and regulation of transposition in addition to dysregulated myelination trajectories following MIA. Moreover, the links between the pathways implicated in our study are ripe for exploration. It will be important to determine if HC-specific changes in neuron projection-related genes drive changes in myelination or vice versa and whether the HC-specific vulnerability in adolescence spreads to other regions with the progression of MIA-induced changes in the brains of offspring. The specific vulnerability of the HC at this age suggests that subcortical pathology may occur earlier than cortical changes, and when coupled with the observed changes in repetitive behavior, warrant examination of striatal and limbic regions in the future. Overall, these results provide a starting point for understanding the molecular changes occurring subsequent to MIA during adolescence, a critical period for development of psychosis.

## Acknowledgments and Disclosures

This project was supported by funding from the Amgen Scholars Foundation awarded to Nicholas Page, an Autism Science Foundation summer undergraduate research grant awarded to Nicholas Page, Michael Gandal, and Daniel Geschwind, and by NIMH grant 5P50MH106438-04 awarded to Daniel Geschwind. This work was supported by a Stanley & Jacqueline Schilling Neuroscience Postdoctoral Research Fellowship (M.L.E.), Dennis Weatherstone Predoctoral Fellowship from Autism Speaks (#7825 M.L.E.), the Letty and James Callinan and Cathy and Andrew Moley Fellowship from the ARCS Foundation (M.L.E.), a Dissertation Year Fellowship from the University of California Office of the President (M.L.E.), P50-MH106438 (C.S.C), and the University of California Davis Research Investments in Science and Engineering Program (A.K.M.).

All authors report no financial or otherwise relevant potential conflicts of interest.

## References

1. Estes ML, McAllister AK (2015): Immune mediators in the brain and peripheral tissues in autism spectrum disorder. Nat Rev Neurosci. 16: 469–486.

2. Hornig M, Bresnahan MA, Che X, Schultz AF, Ukaigwe JE, Eddy ML, et al. (2018): Prenatal fever and autism risk. Mol Psychiatry. 23: 759–766.

3. Brucato M, Ladd-Acosta C, Li M, Caruso D, Hong X, Kaczaniuk J, et al. (2017): Prenatal exposure to fever is associated with autism spectrum disorder in the boston birth cohort. Autism Res. 10: 1878–1890.

4. Sørensen HJ, Mortensen EL, Reinisch JM, Mednick SA (2009): Association between prenatal exposure to bacterial infection and risk of schizophrenia. Schizophr Bull. 35: 631–637.

5. Nielsen PR, Laursen TM, Mortensen PB (2013): Association between parental hospital-treated infection and the risk of schizophrenia in adolescence and early adulthood. Schizophr Bull. 39: 230–237.

6. Khandaker GM, Zimbron J, Lewis G, Jones PB (2013): Prenatal maternal infection, neurodevelopment and adult schizophrenia: a systematic review of population-based studies. Psychological Medicine. 43.

7. Estes M, McAllister A (2016): Maternal immune activation: Implications for neuropsychiatric disorders. Science 353: 772–777.

8. Lichtenstein P, Yip BH, Björk C, Pawitan Y, Cannon TD, Sullivan PF, Hultman CM (2009): Common genetic determinants of schizophrenia and bipolar disorder in Swedish families: a population-based study. Lancet. 373: 234–239.

9. Caspi A, Moffitt T (2006): Gene-environment interactions in psychiatry: joining forces with neuroscience. Nature Reviews Neuroscience 7: 583–590.

10. Tylee D, Sun J, Hess J, Tahir M, Sharma E, Malik R et al. (2018): Genetic correlations among psychiatric and immune-related phenotypes based on genome-wide association data. American Journal of Medical Genetics Part B: Neuropsychiatric Genetics 177: 641–657.

11. Kolevzon A, Gross R, Reichenberg A (2007): Prenatal and Perinatal Risk Factors for Autism. Archives of Pediatrics & Adolescent Medicine 161: 326.

12. Smith SEP, Li J, Garbett K, Mirnics K, Patterson PH (2007): Maternal immune activation alters fetal brain development through interleukin-6. J Neurosci. 27: 10695–10702.

13. Careaga M, Murai T, Bauman MD (2017): Maternal Immune Activation and Autism Spectrum Disorder: From Rodents to Nonhuman and Human Primates. Biol Psychiatry. 81: 391–401.

14. Meyer U (2014): Prenatal Poly(I:C) Exposure and Other Developmental Immune Activation Models in Rodent Systems. Biological Psychiatry 75: 307–315.

15. Knuesel I, Chicha L, Britschgi M, Schobel S, Bodmer M, Hellings J et al. (2014): Maternal immune activation and abnormal brain development across CNS disorders. Nature Reviews Neurology 10: 643–660.

16. Reisinger S, Khan D, Kong E, Berger A, Pollak A, Pollak D (2015): The Poly(I:C)-induced maternal immune activation model in preclinical neuropsychiatric drug discovery. Pharmacology & Therapeutics 149: 213–226.

17. Patterson P (2009): Immune involvement in schizophrenia and autism: Etiology, pathology and animal models. Behavioural Brain Research 204: 313–321.

18. Richetto J, Calabrese F, Riva M, Meyer U (2013): Prenatal Immune Activation Induces Maturation-Dependent Alterations in the Prefrontal GABAergic Transcriptome. Schizophrenia Bulletin 40: 351–361.

19. Hoftman G, Volk D, Bazmi H, Li S, Sampson A, Lewis D (2013): Altered Cortical Expression of GABA-Related Genes in Schizophrenia: Illness Progression vs Developmental Disturbance. Schizophrenia Bulletin 41: 180–191.

20. Schmidt M, Mirnics K (2014): Neurodevelopment, GABA System Dysfunction, and Schizophrenia. Neuropsychopharmacology 40: 190–206.

21. Harvey L, Boksa P (2012): A stereological comparison of GAD67 and reelin expression in the hippocampal stratum oriens of offspring from two mouse models of maternal inflammation during pregnancy. Neuropharmacology 62: 1767–1776.

22. Canetta S, Bolkan S, Padilla-Coreano N, Song L, Sahn R, Harrison N et al. (2016): Maternal immune activation leads to selective functional deficits in offspring parvalbumin interneurons. Molecular Psychiatry 21: 956–968.

23. Wolff A, Bilkey D (2015): Prenatal immune activation alters hippocampal place cell firing characteristics in adult animals. Brain, Behavior, and Immunity 48: 232–243.

24. Zhang Z, van Praag H (2015): Maternal immune activation differentially impacts mature and adult-born hippocampal neurons in male mice. Brain, Behavior, and Immunity 45: 60–70.

25. Bauman MD, Iosif A-M, Smith SEP, Bregere C, Amaral DG, Patterson PH (2014): Activation of the maternal immune system during pregnancy alters behavioral development of rhesus monkey offspring. Biol Psychiatry. 75: 332–341.

26. Machado CJ, Whitaker AM, Smith SEP, Patterson PH, Bauman MD (2015): Maternal immune activation in nonhuman primates alters social attention in juvenile offspring. Biol Psychiatry. 77: 823–832.

27. Bauman MD, Lesh TA, Rowland DJ, Schumann CM, Smucny J, Kukis DL, et al. (2019): Preliminary evidence of increased striatal dopamine in a nonhuman primate model of maternal immune activation. Transl Psychiatry. 9: 135.

28. Parikshak NN, Swarup V, Belgard TG, Irimia M, Ramaswami G, Gandal MJ, et al. (2016): Genome-wide changes in lncRNA, splicing, and regional gene expression patterns in autism. Nature. 540: 423–427.

29. Gandal MJ, Zhang P, Hadjimichael E, Walker RL, Chen C, Liu S, et al. (2018): Transcriptome-wide isoform-level dysregulation in ASD, schizophrenia, and bipolar disorder. Science. 362. doi: 10.1126/science.aat8127.

30. Fillman SG, Cloonan N, Catts VS, Miller LC, Wong J, McCrossin T, et al. (2013): Increased inflammatory markers identified in the dorsolateral prefrontal cortex of individuals with schizophrenia. Mol Psychiatry. 18: 206–214.

31. Voineagu I, Wang X, Johnston P, Lowe JK, Tian Y, Horvath S, et al. (2011): Transcriptomic analysis of autistic brain reveals convergent molecular pathology. Nature. 474: 380–384.

32. Gandal MJ, Haney JR, Parikshak NN, Leppa V, Ramaswami G, Hartl C, et al. (2018): Shared molecular neuropathology across major psychiatric disorders parallels polygenic overlap. Science. 359: 693–697.

33. Volk DW, Chitrapu A, Edelson JR, Roman KM, Moroco AE, Lewis DA (2015): Molecular mechanisms and timing of cortical immune activation in schizophrenia. Am J Psychiatry. 172: 1112–1121.

34. Gupta S, Ellis SE, Ashar FN, Moes A, Bader JS, Zhan J, et al. (2014): Transcriptome analysis reveals dysregulation of innate immune response genes and neuronal activity-dependent genes in autism. Nat Commun. 5: 5748.

35. Ramaker RC, Bowling KM, Lasseigne BN, Hagenauer MH, Hardigan AA, Davis NS, et al. (2017): Post-mortem molecular profiling of three psychiatric disorders. Genome Med. 9: 72.

36. Bray NL, Pimentel H, Melsted P, Pachter L (2016): Near-optimal probabilistic RNA-seq quantification. Nat Biotechnol. 34: 525–527.

37. Robinson MD, McCarthy DJ, Smyth GK (2010): edgeR: a Bioconductor package for differential expression analysis of digital gene expression data. Bioinformatics. 26.

38. Law CW, Chen Y, Shi W, Smyth GK (2014): voom: precision weights unlock linear model analysis tools for RNA-seq read counts. Genome Biol. 15: R29.

39. Garay P, Hsiao E, Patterson P, McAllister A (2013): Maternal immune activation causes age- and region-specific changes in brain cytokines in offspring throughout development. Brain, Behavior, and Immunity 31: 54–68.

40. Langfelder P, Horvath S (2008): WGCNA: an R package for weighted correlation network analysis. BMC Bioinformatics. 9: 559.

41. Criscione SW, Zhang Y, Thompson W, Sedivy JM, Neretti N (2014): Transcriptional landscape of repetitive elements in normal and cancer human cells. BMC Genomics. 15: 583.

42. Winden K, Oldham M, Mirnics K, Ebert P, Swan C, Levitt P et al. (2009): The organization of the transcriptional network in specific neuronal classes. Molecular Systems Biology 5: 291.

43. Reimand J, Arak T, Adler P, Kolberg L, Reisberg S, Peterson H, Vilo J (2016): g:Profiler—a web server for functional interpretation of gene lists (2016 update). Nucleic Acids Res. 44: W83–W89.

44. Skene NG, Grant SGN (2016): Identification of Vulnerable Cell Types in Major Brain Disorders Using Single Cell Transcriptomes and Expression Weighted Cell Type Enrichment. Front Neurosci. 10: 16.

45. Wang D, Liu S, Warrell J, Won H, Shi X, Navarro FCP, et al. (2018): Comprehensive functional genomic resource and integrative model for the human brain. Science. 362. doi: 10.1126/science.aat8464.

46. Plaisier SB, Taschereau R, Wong JA, Graeber TG (2010): Rank-rank hypergeometric overlap: identification of statistically significant overlap between gene-expression signatures. Nucleic Acids Res. 38: e169–e169.

47. Kuramochi-Miyagawa S, Watanabe T, Gotoh K, Totoki Y, Toyoda A, Ikawa M, et al. (2008): DNA methylation of retrotransposon genes is regulated by Piwi family members MILI and MIWI2 in murine fetal testes. Genes Dev. 22: 908–917.

48. Nandi S, Chandramohan D, Fioriti L, Melnick AM, Hébert JM, Mason CE, et al. (2016): Roles for small noncoding RNAs in silencing of retrotransposons in the mammalian brain. Proc Natl Acad Sci U S A. doi: 10.1073/pnas.1609287113.

49. Leighton L, Wei W, Marshall P, Ratnu V, Li X, Zajaczkowski E et al. (2019): Disrupting the hippocampal Piwi pathway enhances contextual fear memory in mice. Neurobiology of Learning and Memory 161: 202–209.

50. Jia L, Liang T, Yu X, Ma C, Zhang S (2013): MGARP Regulates Mouse Neocortical Development via Mitochondrial Positioning. Molecular Neurobiology 49: 1293–1308.

51. Li M, Jaffe AE, Straub RE, Tao R, Shin JH, Wang Y, et al. (2016): A human-specific AS3MT isoform and BORCS7 are molecular risk factors in the 10q24.32 schizophrenia-associated locus. Nat Med. 22: 649–656.

52. Goudarzi S, Rivera A, Butt A, Hafizi S (2016): Gas6 Promotes Oligodendrogenesis and Myelination in the Adult Central Nervous System and After Lysolecithin-Induced Demyelination. ASN Neuro 8: 175909141666843.

53. Funakoshi H, Yonemasu T, Nakano T, Matumoto K, Nakamura T (2002): Identification of Gas6, a putative ligand for Sky and Axl receptor tyrosine kinases, as a novel neurotrophic factor for hippocampal neurons. Journal of Neuroscience Research 68: 150–160.

54. Miyata M, Mandai K, Maruo T, Sato J, Shiotani H, Kaito A et al. (2016): Localization of nectin-2δ at perivascular astrocytic endfoot processes and degeneration of astrocytes and neurons in nectin-2 knockout mouse brain. Brain Research 1649: 90–101.

55. Workman AD, Charvet CJ, Clancy B, Darlington RB, Finlay BL (2013): Modeling transformations of neurodevelopmental sequences across mammalian species. J Neurosci. 33: 7368–7383.

56. Amini-Bavil-Olyaee S, Choi YJ, Lee JH, Shi M, Huang I-C, Farzan M, Jung JU (2013): The antiviral effector IFITM3 disrupts intracellular cholesterol homeostasis to block viral entry. Cell Host Microbe. 13: 452–464.

57. Ibi D, Nagai T, Nakajima A, Mizoguchi H, Kawase T, Tsuboi D, et al. (2013): Astroglial IFITM3 mediates neuronal impairments following neonatal immune challenge in mice. Glia. 61: 679–693.

58. Mi H, Muruganujan A, Ebert D, Huang X, Thomas P (2018): PANTHER version 14: more genomes, a new PANTHER GO-slim and improvements in enrichment analysis tools. Nucleic Acids Research 47: D419–D426.

59. Mager DL, Hunter DG, Schertzer M, Freeman JD (1999): Endogenous retroviruses provide the primary polyadenylation signal for two new human genes (HHLA2 and HHLA3). Genomics. 59: 255–263.

60. Ito J, Sugimoto R, Nakaoka H, Yamada S, Kimura T, Hayano T, Inoue I (2017): Systematic identification and characterization of regulatory elements derived from human endogenous retroviruses. PLoS Genet. 13: e1006883.

61. Tam O, Ostrow L, Gale Hammell M (2019): Diseases of the nERVous system: retrotransposon activity in neurodegenerative disease. Mobile DNA 10:. doi: 10.1186/s13100-019-0176-1.

62. Gardner E, Prigmore E, Gallone G, Danecek P, Samocha K, Handsaker J et al. (2019): Contribution of retrotransposition to developmental disorders. Nature Communications 10:. doi: 10.1038/s41467-019-12520-y.

63. Nave K-A, Werner HB (2014): Myelination of the nervous system: mechanisms and functions. Annu Rev Cell Dev Biol. 30: 503–533.

64. Vanier MT (2010): Niemann-Pick disease type C. Orphanet J Rare Dis. 5: 16.

65. Fischer von Mollard G, Stahl B, Khokhlatchev A, Südhof TC, Jahn R (1994): Rab3C is a synaptic vesicle protein that dissociates from synaptic vesicles after stimulation of exocytosis. Journal of Biological Chemistry 269: 10971–10974.

66. Pavlos N, Gronborg M, Riedel D, Chua J, Boyken J, Kloepper T et al. (2010): Quantitative Analysis of Synaptic Vesicle Rabs Uncovers Distinct Yet Overlapping Roles for Rab3a and Rab27b in Ca2+-Triggered Exocytosis. Journal of Neuroscience 30: 13441–13453.

67. Renden R, Berwin B, Davis W, Ann K, Chin C, Kreber R et al. (2001): Drosophila CAPS Is an Essential Gene that Regulates Dense-Core Vesicle Release and Synaptic Vesicle Fusion. Neuron 31: 421–437.

68. Giovanoli S, Notter T, Richetto J, Labouesse MA, Vuillermot S, Riva MA, Meyer U (2015): Late prenatal immune activation causes hippocampal deficits in the absence of persistent inflammation across aging. J Neuroinflammation. 12: 221.

69. Notter T, Coughlin J, Gschwind T, Weber-Stadlbauer U, Wang Y, Kassiou M et al. (2017): Translational evaluation of translocator protein as a marker of neuroinflammation in schizophrenia. Molecular Psychiatry 23: 323–334.

70. Richetto J, Massart R, Weber-Stadlbauer U, Szyf M, Riva MA, Meyer U (2017): Genome-wide DNA Methylation Changes in a Mouse Model of Infection-Mediated Neurodevelopmental Disorders. Biol Psychiatry. 81: 265–276.

71. Labouesse MA, Dong E, Grayson DR, Guidotti A, Meyer U (2015): Maternal immune activation induces GAD1 and GAD2 promoter remodeling in the offspring prefrontal cortex. Epigenetics. 10: 1143–1155.

72. Basil P, Li Q, Dempster EL, Mill J, Sham P-C, Wong CCY, McAlonan GM (2014): Prenatal maternal immune activation causes epigenetic differences in adolescent mouse brain. Transl Psychiatry. 4: e434.

73. Connor CM, Dincer A, Straubhaar J, Galler JR, Houston IB, Akbarian S (2012): Maternal immune activation alters behavior in adult offspring, with subtle changes in the cortical transcriptome and epigenome. Schizophr Res. 140: 175–184.

74. Bundo M, Toyoshima M, Okada Y, Akamatsu W, Ueda J, Nemoto-Miyauchi T, et al. (2014): Increased l1 retrotransposition in the neuronal genome in schizophrenia. Neuron. 81: 306–313.

75. Suarez N, Macia A, Muotri A (2017): LINE-1 retrotransposons in healthy and diseased human brain. Developmental Neurobiology 78: 434–455.

76. Erwin J, Marchetto M, Gage F (2014): Mobile DNA elements in the generation of diversity and complexity in the brain. Nature Reviews Neuroscience 15: 497–506.

77. Davis K, Stewart D, Friedman J, Buchsbaum M, Harvey P, Hof P et al. (2003): White Matter Changes in Schizophrenia. Archives of General Psychiatry 60: 443.

78. Scheel M, Prokscha T, Bayerl M, Gallinat J, Montag C (2012): Myelination deficits in schizophrenia: evidence from diffusion tensor imaging. Brain Structure and Function 218: 151–156.

79. Makinodan M, Tatsumi K, Manabe T, Yamauchi T, Makinodan E, Matsuyoshi H, et al. (2008): Maternal immune activation in mice delays myelination and axonal development in the hippocampus of the offspring. J Neurosci Res. 86: 2190–2200.

80. Antonson AM, Balakrishnan B, Radlowski EC, Petr G, Johnson RW (2018): Altered Hippocampal Gene Expression and Morphology in Fetal Piglets following Maternal Respiratory Viral Infection. Dev Neurosci. 40: 104–119.

81. Hakak Y, Walker J, Li C, Wong W, Davis K, Buxbaum J et al. (2001): Genome-wide expression analysis reveals dysregulation of myelination-related genes in chronic schizophrenia. Proceedings of the National Academy of Sciences 98: 4746–4751.

82. Graciarena M, Seiffe A, Nait-Oumesmar B, Depino A (2019): Hypomyelination and Oligodendroglial Alterations in a Mouse Model of Autism Spectrum Disorder. Frontiers in Cellular Neuroscience 12:. doi: 10.3389/fncel.2018.00517.

83. Kentner AC, Bilbo SD, Brown AS, Hsiao EY, McAllister AK, Meyer U, et al. (2018): Maternal immune activation: reporting guidelines to improve the rigor, reproducibility, and transparency of the model. Neuropsychopharmacology. doi: 10.1038/s41386-018-0185-7.

84. Meyer U (2019): Neurodevelopmental Resilience and Susceptibility to Maternal Immune Activation. Trends Neurosci. doi: 10.1016/j.tins.2019.08.001.

85. Garbett KA, Hsiao EY, Kálmán S, Patterson PH, Mirnics K (2012): Effects of maternal immune activation on gene expression patterns in the fetal brain. Transl Psychiatry. 2: e98.

86. Lombardo MV, Moon HM, Su J, Palmer TD, Courchesne E, Pramparo T (2018): Maternal immune activation dysregulation of the fetal brain transcriptome and relevance to the pathophysiology of autism spectrum disorder. Mol Psychiatry. 23: 1001–1013.

87. Oskvig DB, Elkahloun AG, Johnson KR, Phillips TM, Herkenham M (2012): Maternal immune activation by LPS selectively alters specific gene expression profiles of interneuron migration and oxidative stress in the fetus without triggering a fetal immune response. Brain Behav Immun. 26: 623–634.

88. Weber-Stadlbauer U, Richetto J, Labouesse MA, Bohacek J, Mansuy IM, Meyer U (2017): Transgenerational transmission and modification of pathological traits induced by prenatal immune activation. Mol Psychiatry. 22: 102–112.

89. Farrelly L, Föcking M, Piontkewitz Y, Dicker P, English J, Wynne K, et al. (2015): Maternal immune activation induces changes in myelin and metabolic proteins, some of which can be prevented with risperidone in adolescence. Dev Neurosci. 37: 43–55.

90. Zoghbi H, Bear M (2012): Synaptic Dysfunction in Neurodevelopmental Disorders Associated with Autism and Intellectual Disabilities. Cold Spring Harbor Perspectives in Biology 4: a009886–a009886.

91. Bourgeron T (2015): From the genetic architecture to synaptic plasticity in autism spectrum disorder. Nature Reviews Neuroscience 16: 551–563.

92. Rodenas-Cuadrado P, Ho J, Vernes S (2013): Shining a light on CNTNAP2: complex functions to complex disorders. European Journal of Human Genetics 22: 171–178.

93. McAllister A, Lo D, Katz L (1995): Neurotrophins regulate dendritic growth in developing visual cortex. Neuron 15: 791–803.

94. Xiao J, Wong AW, Willingham MM, van den Buuse M, Kilpatrick TJ, Murray SS (2010): Brain-derived neurotrophic factor promotes central nervous system myelination via a direct effect upon oligodendrocytes. Neurosignals. 18: 186–202.

95. Collins C, Airey D, Young N, Leitch D, Kaas J (2010): Neuron densities vary across and within cortical areas in primates. Proceedings of the National Academy of Sciences 107: 15927–15932.

96. Militerni R, Bravaccio C, Falco C, Fico C, Palermo M (2002): European Child & Adolescent Psychiatry 11: 210–218.

97. Langen M, Bos D, Noordermeer S, Nederveen H, van Engeland H, Durston S (2014): Changes in the Development of Striatum Are Involved in Repetitive Behavior in Autism. Biological Psychiatry 76: 405–411.

98. Lieberman J, Small S, Girgis R (2019): Early Detection and Preventive Intervention in Schizophrenia: From Fantasy to Reality. American Journal of Psychiatry 176: 794–810.

