## Supplemental Figures for "Alterations in retrotransposition, synaptic connectivity, and myelination implicated by transcriptomic changes following maternal immune activation in non-human primates"

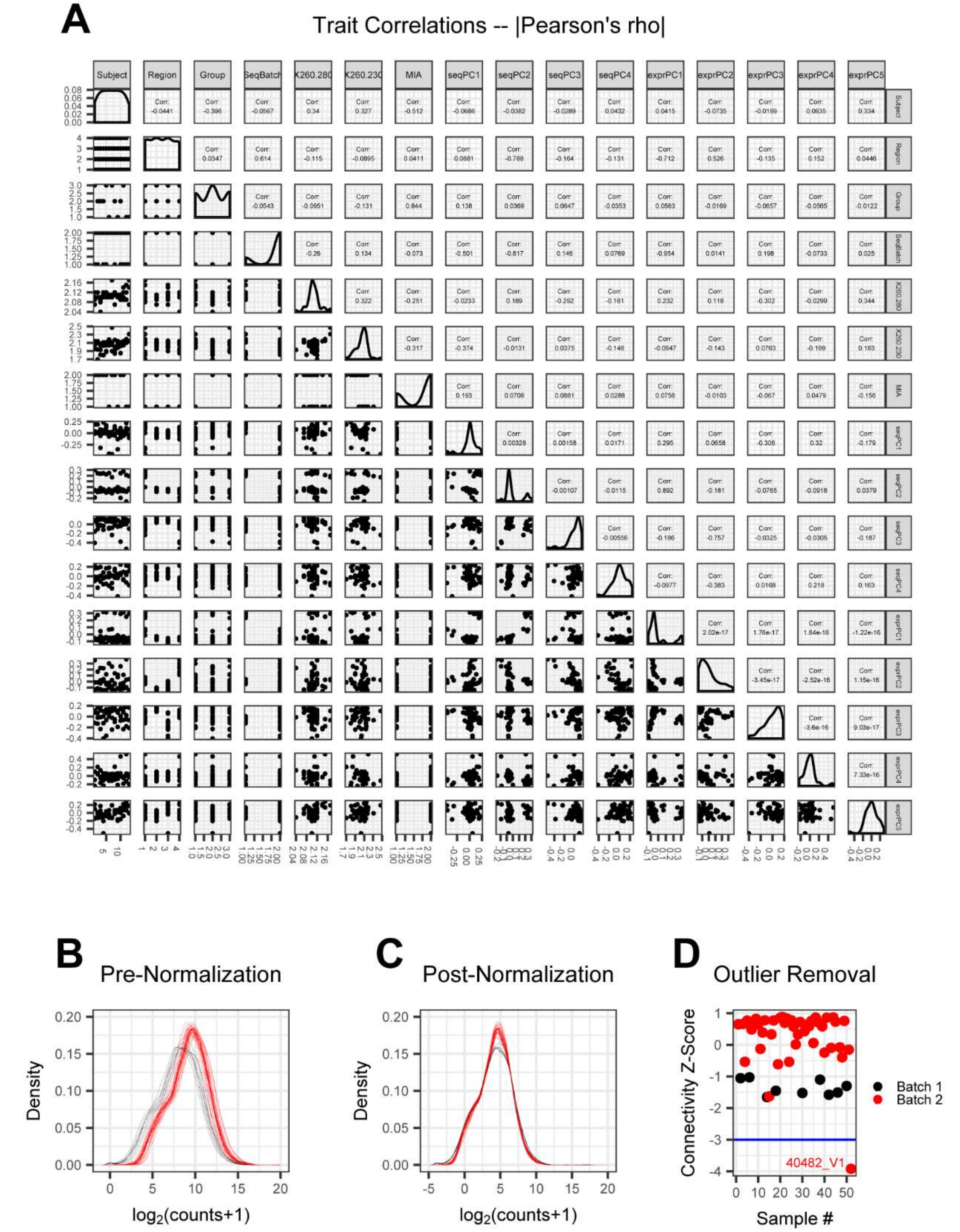


(Figure S1)

**Figure S1.** Quality control for RNA-seq analysis. **(A)** Correlations between all experimental covariates were used to determine the number of sequencing principle components to correct for during differential expression analysis. High correlations between SeqPCs 1-4 and other technical co-variates indicate that the majority of technical variation in our analysis is captured by the SeqPCs. **(B, C)** Read count density across all samples before and after TMM normalization. **(D)** All samples with a connectivity z-score less than -3 were considered outliers and were removed from all further analysis.

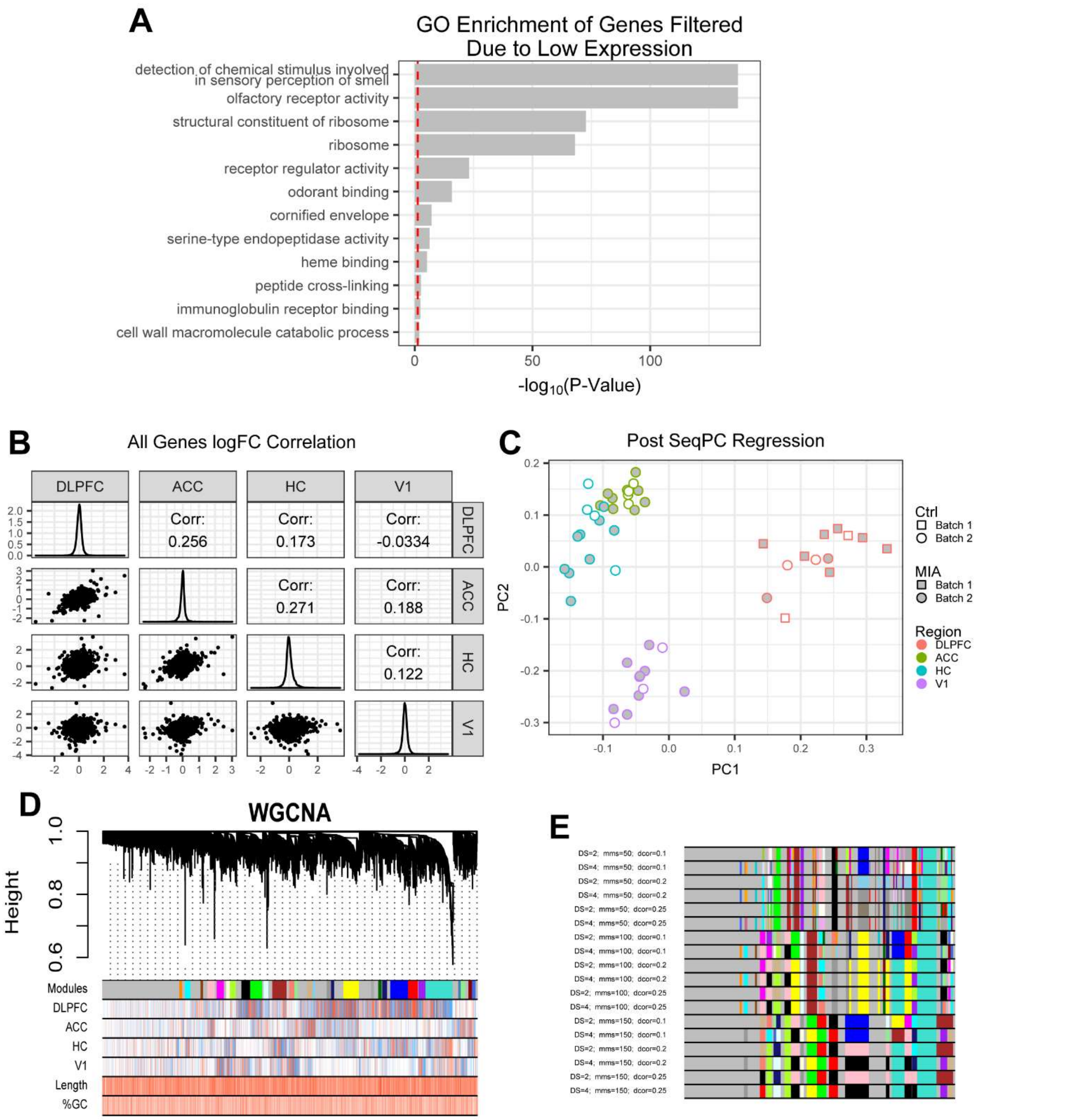


(Figure S2)

**Figure S2.** Technical analyses related to Figures 1-4. **(A)** All significantly enriched GO terms among the 6,410 low or unexpressed genes removed from downstream analysis. Red dotted line indicates an FDR significance threshold of 0.05. **(B)** $log_2$(FC) of all genes are compared pairwise with each other brain region. Small correlations indicated minimal overlap in the  similarity of gene expression changes between each region following MIA. **(C)** Top expression principle components across all samples after regressing out the variance explained by SeqPCs1-4. **(D)** WGCNA was used to construct a signed bicor network and identify co-expression network modules, each containing genes whose expression is highly correlated across samples. Dendrotree height indicates the degree of co-expression correlation, colors indicate co-expression network module assignments. Colors also indicate the relative expression of each gene within a given brain region (top) followed by a heatmap of gene length and GC-content (bottom). **(E)** Iteration through multiple dendrotree cut setting for WGCNA.

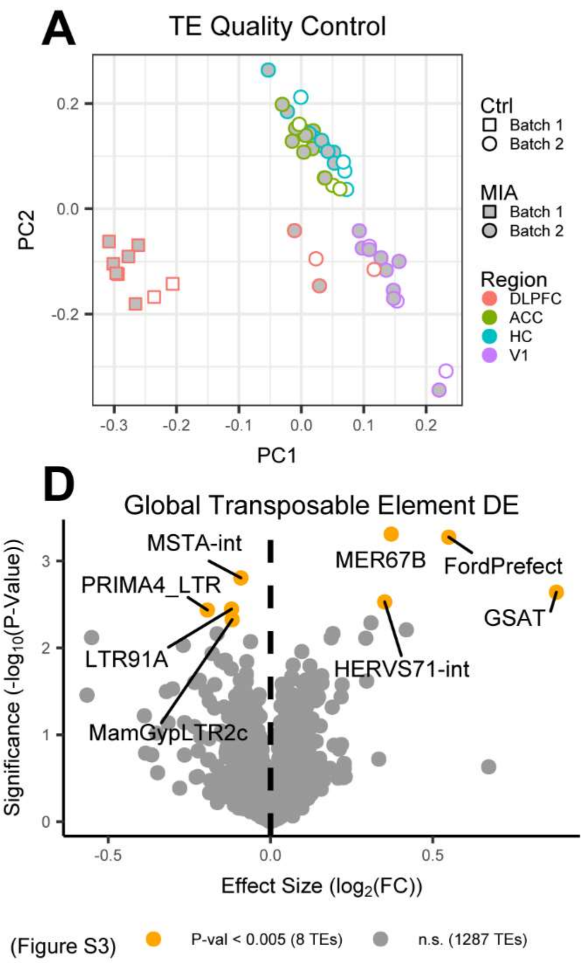


**Figure S3.** Quality control for transposable elements analysis. **(A)** Top principal components (PCs) of normalized transposable element expression demonstrating that batch and brain region are the largest drivers of variation in the dataset. **(B, C)** Transposable element read count density across all samples before and after TMM normalization. **(D)** Differential transposable element expression analysis using Limma-voom was performed on all saline injected vs. poly:ICLC injected MIA samples pooled across brain regions and MIA timepoints. Volcano plots indicate all transposable elements with suggestive association with MIA in each brain region. There are no transposable elements that pass FDR correction for differential transposable elements expression (FDR<0.1). Yellow dots indicate suggestive association with MIA (p<0.005) and grey dots indicate minimal or no association. **(E)** Selected boxplot of *PiggyBac Transposable Element Derived 3* (*PGBD3*) expression in the cingulate cortex. **(F)** Selected boxplot of *PiggyBac Transposable Element Derived 5* (*PGBD5*) expression in the hippocampus.

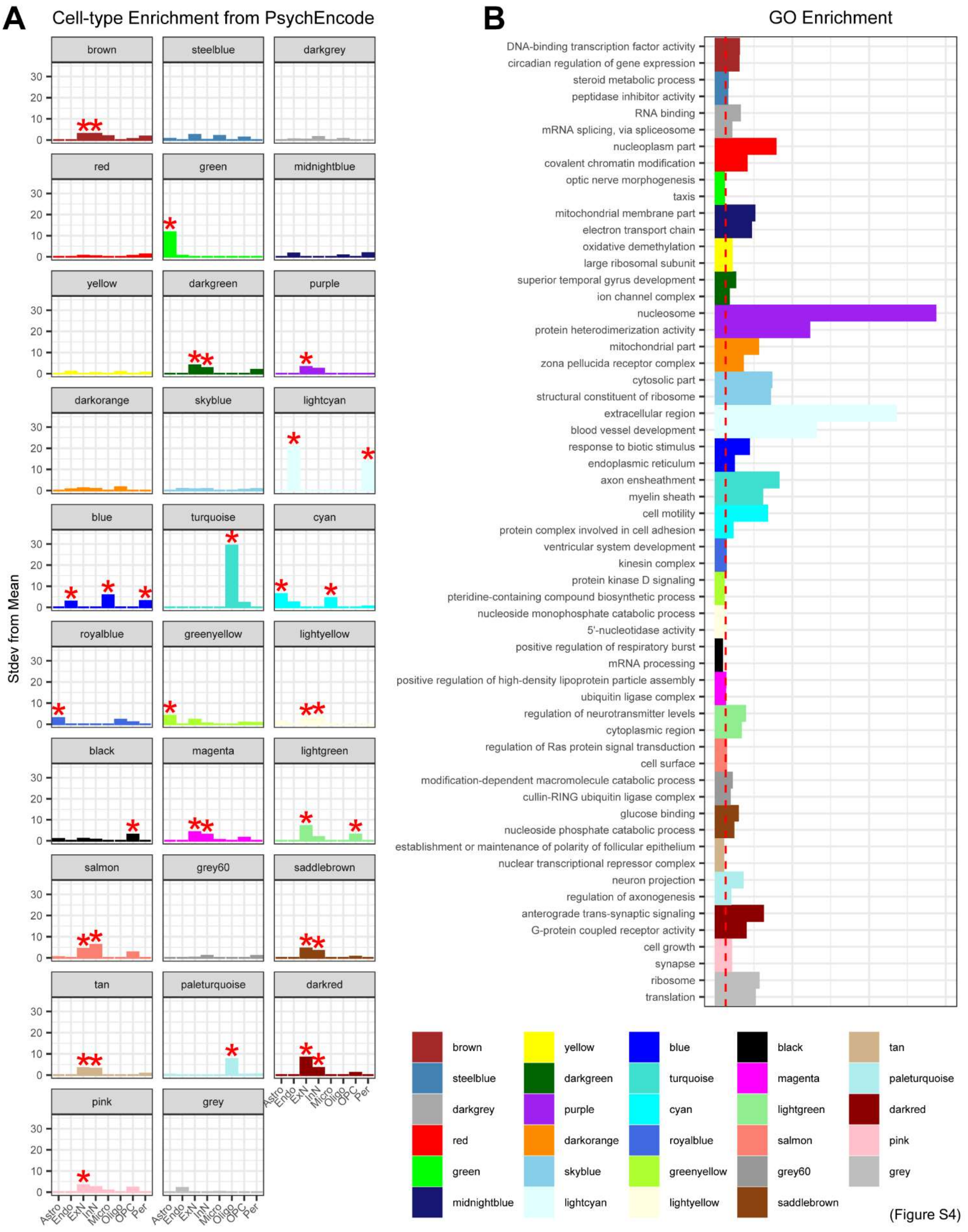


**Figure S4.** Enrichment analysis related to WGCNA. **(A)** Cell-type specificity of all co-expression network modules following MIA based on PsychEncode and Lake et al., 2018 adult human single-cell nuc-Seq. *FDR<0.05, unlabeled cell-types are not significantly enriched. **(B)** Top two GO terms enriched among co-expression network modules following MIA determined using g:ProfileR. Red dotted line indicates an FDR significance threshold of 0.05.

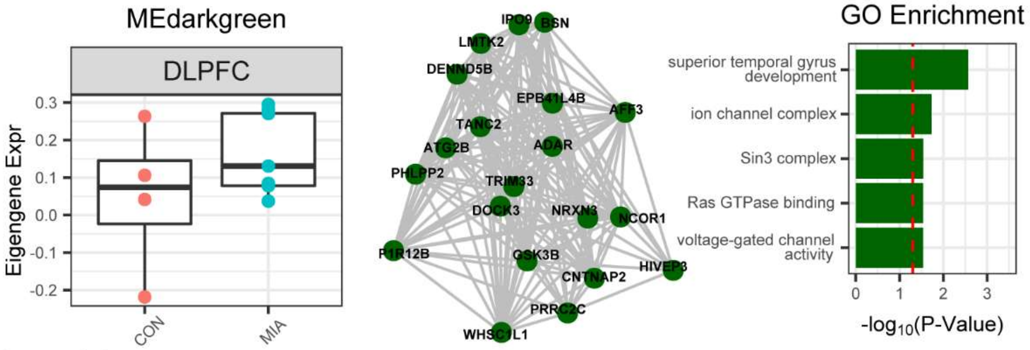


(Figure S5)

**Figure S5.** Transcriptomic differences in the DLPFC implicate development and ion channels. **(A)** Boxplot of MEdarkgreen module eigengene expression in the DLFPC. **(B)** Top 20 hub genes for MEdarkgreen. **(C)** Top GO terms enriched in MEdarkgreen region by g:ProfileR. For all GO enrichment plots, red dotted line indicates an FDR significance threshold of 0.05.

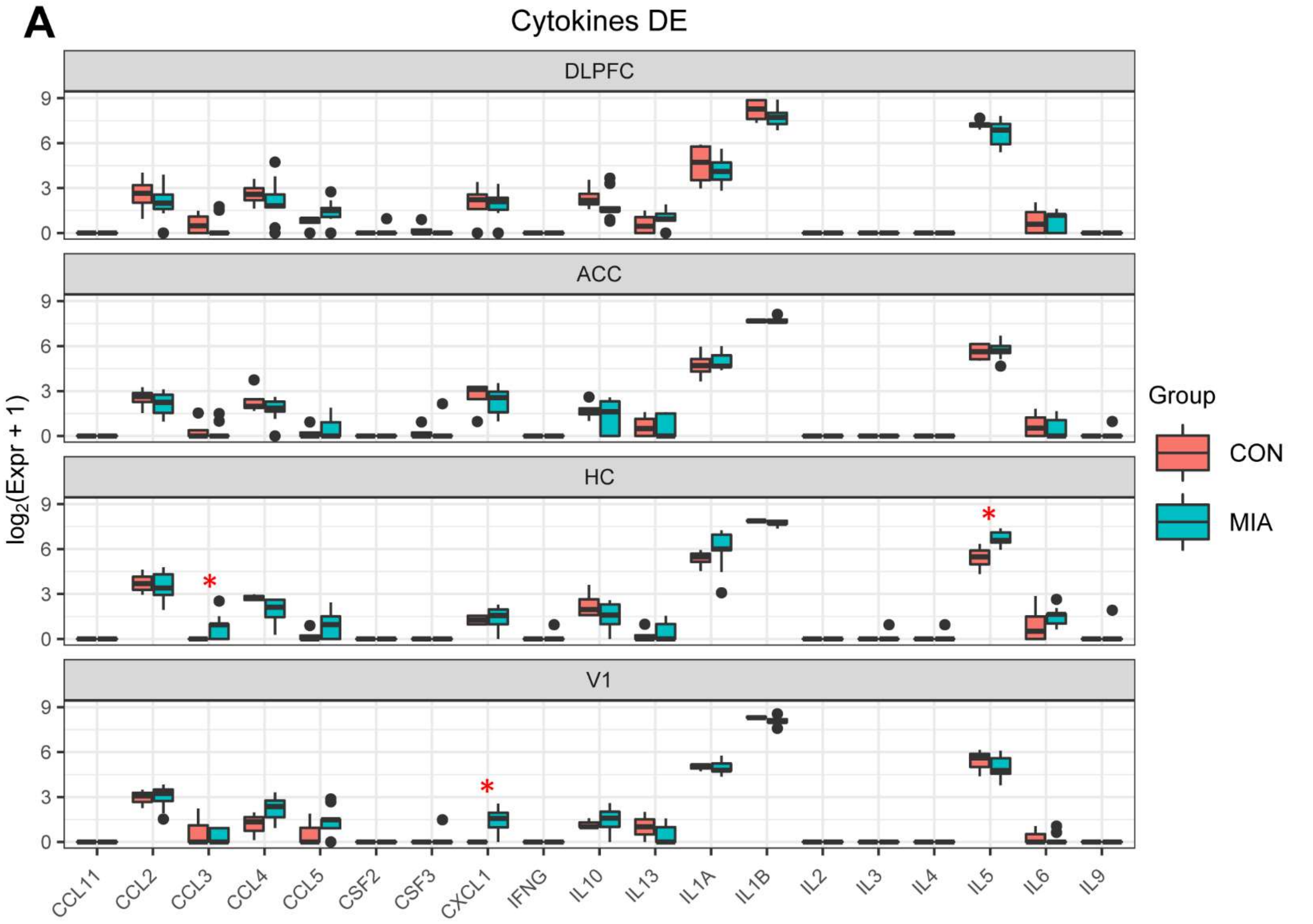


(Figure S6)

**Figure S6.** Specific changes in cytokine expression following MIA. **(A)** Differential cytokine expression was determined using Limma-lmFit separately for each brain region on all saline injected vs. poly:ICLC injected MIA samples pooled across MIA timepoints. Analysis was performed on all 19 annotated cytokines in the Rhesus genome regardless of low-expressed gene filtering criteria applied in other analyses. *p<0.05.

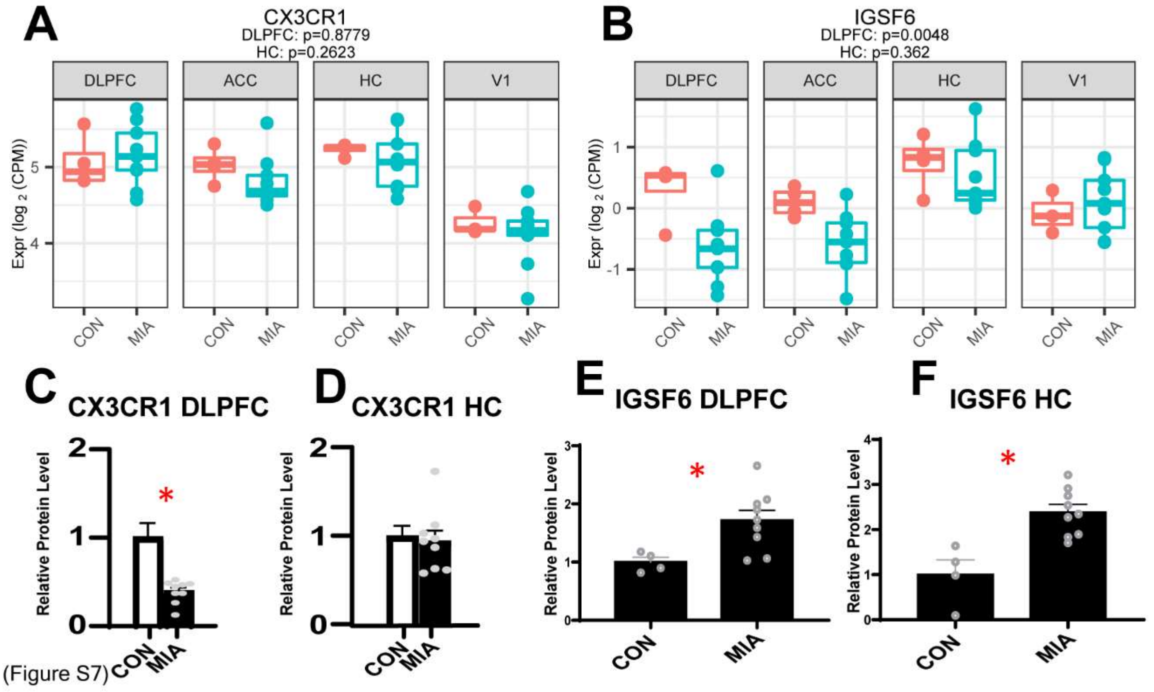


**Figure S7.** Contrasting mRNA and protein expression changes of microglia cell-type markers. **(A, B)** Boxplots of RNA-seq gene expression results for microglial cell-type markers *CX3CR1* and *IGSF6* respectively. Western blot quantifications of CX3CR1 **(C, D)** and IGSF6 **(E, F)** protein in DLPFC and HC respectively following MIA. *****$p<0.05$.

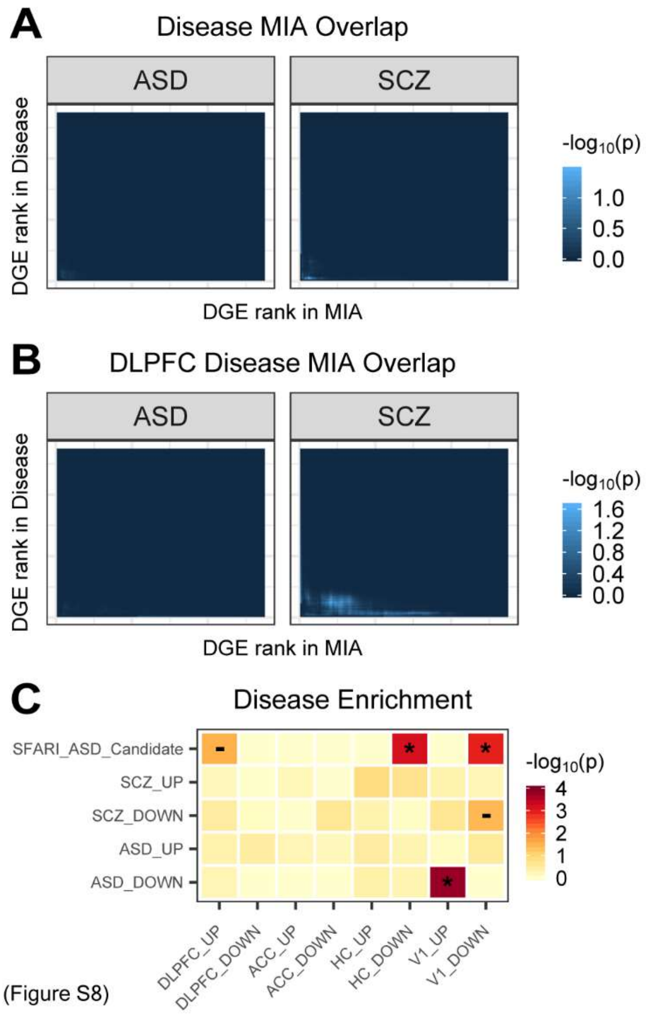


**Figure S8.** Cross disorder overlaps with transcriptomic changes following MIA. **(A)** RRHO between disease differentially expressed ranked gene lists from the PsychEncode consortium and the global differential expression analysis from this study (Figure 2A). **(B)** RRHO between disease differentially expressed ranked gene lists from the PsychEncode consortium and the DLPFC differential expression analysis from this study (Figure 2A). **(C)** Enrichment of disease associated gene lists with regional up and down regulated gene lists from this study. * indicates enrichment with Bonferroni corrected FDR < 0.05. – indicates enrichment with uncorrected p < 0.05.

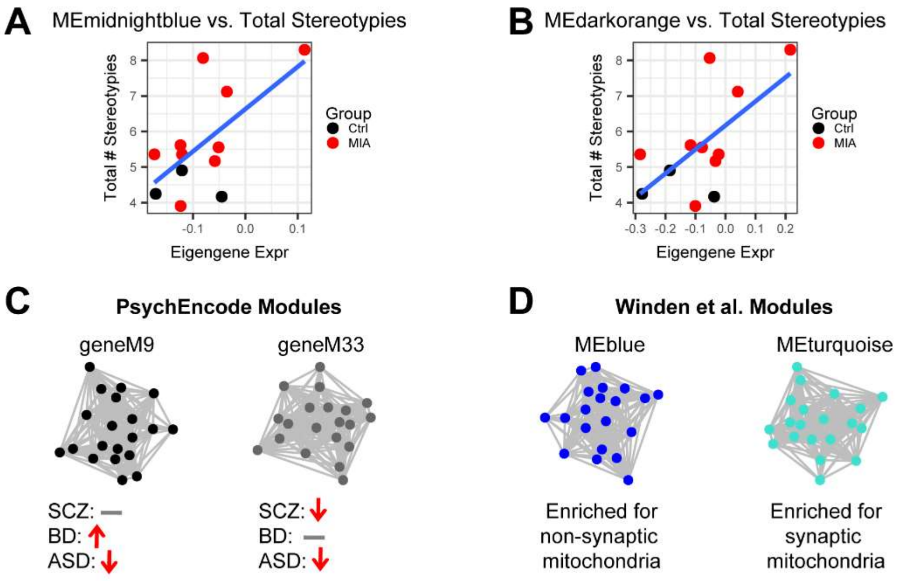


| Monkey MIA | PsychEncode | Fishers P-val | FDR | Overlap |
|---|---|---|---|---|
| MEmidnightblue | geneM9 | 0.89 | 1 | PSMC6, NDUFV1, ZFYVE27 |
| MEmidnightblue | geneM33 | 4.7e-6 | 0.005 | SLIRP, USMG5, COX7C, C14orf2, PET100, SNRPD2, COX7B, COX6C |
| MEdarkorange | geneM9 | 0.59 | 1 | RAN, PGAM1 |
| MEdarkorange | geneM33 | 3.7e-5 | 0.04 | HINT1, COX6B1, ATP5I, ATP5G1, UQCRH |

| Monkey MIA | Winden et al. 2009 | Fishers P-val | FDR | Overlap |
|---|---|---|---|---|
| MEmidnightblue | MEblue | 0.70 | 1 | AUH |
| MEmidnightblue | MEblue | 0.03 | 1 | ANK3, MTX2, PGAM1, RAP1G, DS1, MTCH2 |
| MEdarkorange | MEturquoise | 0.69 | 1 | ARHGAP5, MKKS |
| MEdarkorange | MEturquoise | 0.11 | 1 | ATP5F1, GFM1, PDCD10, COPS2, UBE2D1, RAD21, BZW1 |

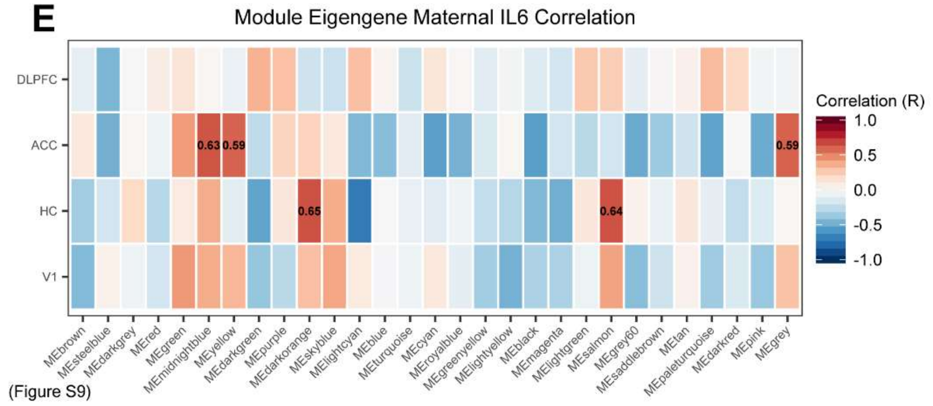


(Figure S9)

**Figure S9.** Correlation between maternal IL-6, behavioral measurements, and module eigengenes. **(A)** Plot of $\log_2$(Total # stereotypies) versus MEmidnightblue module eigengene expression. **(B)** Plot of $\log_2$(Total # stereotypies) versus MEdarkorange module eigengene expression. **(C)** Overlap between MEmidnightblue, MEdarkorange, and mitochondria enriched modules from PsychEncode. Red arrows indicate significant up or down-regulation of PsychEncode modules in human post-mortem brain tissue across disorders, p-values are reported as determined via a one-sided Fisher's exact test. FDR corrections were performed on all p-values determined from pairwise over-enrichment analysis between all NHP MIA and PsychEncode modules. **(D)** Overlap between MEmidnightblue, MEdarkorange, and cellular compartment specific mitochondria enriched modules from Winden et al. (ref (42)). Text indicates whether mitochondrial modules from Winden et al are enriched for synaptic or non-synaptic genes. p-values are reported as determined via a one-sided Fisher's exact test. FDR corrections were performed on all p-values determined from pairwise over-enrichment analysis between all NHP MIA and Winden et al. modules. **(E)** Correlation between module eigengene expression in a given region and the $\log_2$(maternal IL-6 levels) in a given subject.

### Supplementary Tables

**Table S1.** Sample metadata and SeqPCs.

**Table S2.** Global and regional differential gene expression results.

**Table S3.** 1st and 2nd trimester specific differential gene expression results.

**Table S4.** Co-expression network membership and kME for each gene determined by WGCNA.

**Table S5.** Maternal IL6 and behavioral measurements from NHPs.
